# Targeting therapeutic nanoparticles to the glioblastoma resection margin by harnessing post-operative spatiotemporal blood-brain barrier disruption

**DOI:** 10.1101/2025.03.29.646102

**Authors:** Lorena F Fernandes, Chariya Peeyatu, Ben R Dickie, Yu Siong Ho, Lydia A Thompson, Noelia Hernandez, Neus Lozano, Kostas Kostarelos, Thomas Kisby

**Affiliations:** Centre for Nanotechnology in Medicine, School of Biological Sciences, Faculty of Biology, Medicine and Health, University of Manchester, Manchester, UK; Geoffrey Jefferson Brain Research Centre, Manchester Academic Health Science Centre, Northern Care Alliance NHS Foundation Trust, University of Manchester, Manchester, United Kingdom; Division of Informatics, Imaging and Data Sciences, School of Health Sciences, University of Manchester, Manchester, UK; Nanomedicine Lab, Catalan Institute of Nanoscience and Nanotechnology (ICN2), CSIC and BIST, Campus UAB, 08193 Barcelona, Spain; Departament de Química, Facultat de Ciències, Universitat Autònoma de Barcelona, 08193, Bellaterra, Spain; Institute of Neuroscience, Universitat Autònoma de Barcelona, 08913 Barcelona, Spain; Institució Catalana de Recerca i Estudis Avançats (ICREA), Pg. Lluís Companys 23, Barcelona, Spain

## Abstract

Resection surgery is the first-line therapy for glioblastoma (GBM) that is performed in >70% of patients, typically within days of suspected diagnosis. Current protocols for follow-on chemoradiotherapy have shown only modest efficacy in eliminating residual disease, leading to inevitable tumour recurrence. There remains a need for new approaches to swiftly and effectively treat post-operative residual disease to prevent the rapid early progression of recurrent GBM. Using syngeneic preclinical models of glioblastoma resection, we identified a spatially and temporally restricted window of blood brain barrier (BBB) disruption localised to the resection margin, during the immediate (15 min) and early (48-72h) postoperative periods. Intravenous administration of fluorescently labelled, clinically-used liposome nanoparticles during these periods demonstrated that selective accumulation at the postoperative resection margin, while largely being excluded from areas of the brain with an intact BBB, could be achieved. Confocal analysis confirmed the presence of extravasated nanoparticles within the margin parenchyma which largely interacted with microglial populations closely associated with residual tumour cells. Exploiting this, we performed intravenous administration of doxorubicin-loaded liposomes (DOX-Lipo) coinciding with the peak of postoperative BBB disruption and demonstrated both enhanced chemotherapy delivery and consequently complete inhibition of tumour recurrence from a single administration. Overall, this work underscores the importance of timing concomitant chemotherapy to the post-operative timeframe and demonstrates that clinically-used liposomal nanomedicines could be readily repurposed for early post-operative therapy in aggressive brain tumours.

## Introduction

Glioblastoma (GBM) is the most common, aggressive, and neurologically destructive brain cancer in adults^1^. Maximal safe resection in the form of a gross total, or supratotal resection is the first step in treatment, however, due the highly invasive properties of GBM, residual cancer cells will always remain after surgery^2, 3^. The current standard of care (SoC; aka the Stupp protocol) aims to address this with a combined approach of radiotherapy and concurrent/adjuvant temozolomide (TMZ) chemotherapy^4^. Despite this combinatorial approach, recurrence is largely inevitable with current median survival remaining around 15 months post diagnosis and 5 year survival of less than 10%^1, 5^. Notably, in the current SoC, there is a period between resection surgery and the start of chemoradiotherapy of around 4-6 weeks where no anti-tumour therapy is given. During this period, which exceeds the volumetric doubling time of the disease^6, 7^, the presence of early recurrent lesions in the form of rapid early progression (REP) has been identified in up to 50% of patients before the start of chemoradiation^8, 9^. Moreover, this REP has a significant negative impact on prognosis highlighting the potential critical nature of this treatment gap in the outcomes of patients.

Due to the intracranial location of GBM, the blood brain barrier (BBB) is considered a significant limiting factor to the delivery and efficacy of systemically administered follow-on therapy. The BBB is a dynamic barrier composed of endothelial cells connected by relatively impermeable tight junctions further reinforced by astrocyte foot processes surrounding the vessels, and pericytes influencing its functionality^10^. Brain endothelial cells also express a variety of efflux transporters further limiting drug concentrations in the brain^11, 12^. While GBM is largely characterized by a dysfunctional and leaky BBB, this is heterogeneous and is restricted to the bulk of the tumour with peritumoral BBB remaining intact. Indeed, the gadolinium enhanced region as seen in dynamic contrast enhanced (DCE)-MRI (a classical imaging biomarker of BBB leakiness) is used to delineate the target for gross total resection. The effect of this surgery on the remaining BBB is currently unknown, however at the time of follow-on therapy (4-6 weeks after surgery) residual invasive tumour cells that escape resection appear protected by the BBB which restricts effective drug delivery to these cell populations. Consistent with this, postoperative tumour recurrence typically occurs within the resection margin (1-2 cm from the cavity edge) highlighting this site as a key target for GBM therapeutics^2, 3, 13, 14^.

Various nanoparticles have demonstrated an ability to promote selective or targeted drug delivery to tumours in addition to offering improved safety when considering delivery of chemotherapeutics^15, 16^. In the context of brain delivery, intricate approaches to design nanoparticles that can cross the BBB such as increasing interactions with endothelial transporters or hijacking immune cells have been described with varying levels of success^17–19^. At the clinical level, liposomes, which are nanosized spherical vesicles composed of a phospholipid bilayer, have the longest history of use with longstanding FDA/EMA approvals for various indications^16^. The first clinically approved liposome formulation for cancer therapy was liposomal doxorubicin (Doxil^®^/Caelyx^®^) which continues to be applied in the treatment of various solid tumours due to its excellent biodistribution and safety profile compared to standard doxorubicin HCl (Adriamycin). However, due to their relatively large size (80-120 nm) and surface chemistry, liposome formulations such as Doxil^®^ do not readily cross the BBB which has limited their application in neurological and neuro-oncological applications. In contrast, liposomes have been shown to selectively accumulate in the brain in response to brain injuries such as stroke^20, 21^ and traumatic brain injury^22, 23^, where gross BBB dysfunction is known to occur. In both these examples the accumulation of liposomes is both transient or has a biphasic kinetic profile related to the pathophysiology of BBB disruption and is spatially restricted to the sites of injury. Based on this, we hypothesised that the ‘injury’ that occurs as part of surgical resection of GBM may disrupt the local BBB, facilitating local translocation of circulating liposomes into the resection margin.

Here we employed the clinically approved Doxil^®^/Caelyx^®^ formulation to investigate the translocation of liposomes into the brain following resection surgery using preclinical models of GBM resection. We demonstrate selective accumulation of intravenously administered liposomes within the resection margin of GBM when injected at two specific time periods post-resection. Such localization could be exploited for early postoperative therapy and indeed we applied the clinically-used liposomal doxorubicin formulation and demonstrated enhanced efficacy in preventing tumour recurrence. Early post-operative administration within these temporal windows of BBB disruption was essential for improved therapeutic efficacy and highlights the importance of considering such critical time windows in the SoC when developing new therapeutic approaches.

## Results

### Selective accumulation of intravenously administered liposomes in the postoperative resection margin

Based on the GBM clinical journey, maximal safe resection surgery is the standard protocol applied to most (>70%) patients^2, 24^. To better model this we first established a murine model of GBM resection whereby surgical removal of an intracranial cortical tumour was performed by craniotomy and tissue aspiration, mimicking the clinical procedure (**Fig. S1**). Resection of an established GL261-luciferase tumour (7 days after inoculation) effectively eliminated the tumour bioluminescence signal detectable by *in vivo* optical imaging 2 days and 7 days after resection (**Fig. S1a**). Consistent with this, histological assessment evidenced the absence of tumour bulk consistent with the clinical scenario (**Fig. S1b**). Longitudinal monitoring confirmed the eventual recurrence of GBM originating from the resection margin (**Fig. S1c-d**). This surgical approach provided a 2.3 fold increase in median survival (40 days) compared to unresected mice (17.5 days), which is consistent with the relative fold-change survival benefit achieved from gross total resection in patients^24^.

To assess the possible translocation of nanoparticles into the brain following resection surgery we used empty DiI labelled pegylated liposomes based on the clinically approved Doxil^®^/Caelyx^®^ formulation (HSPC:Chol:DSPE-PEG_2000_) (**Fig. S2**). This formulation is clinically approved for various solid extracranial tumours but does not typically enter the brain in the absence of BBB disruption^20, 25^. A single intravenous injection of empty DiI-liposomes was performed at different timepoints either before resection surgery (-15’) or after (+15’, 3h, 24h, 48h, 72h, 7 days) (**Fig. 1a**). The post-surgery injections were timed to ensure that the DiI-liposomes were administered only after stabilization of bleeding and wound closure. Animals were sacrificed 24 hours after the administration of DiI-liposomes and perfused to remove any liposomes remaining in the circulation before DiI fluorescence in organs was measured *ex vivo* (**Fig. 1b** and **Fig. S3**). Selective accumulation of DiI-liposomes was observed around the resection site (right hemisphere) of the perfused brains compared to an absence of fluorescence signal on the contralateral, non-surgically manipulated site (**Fig. 1b**). Quantification of DiI fluorescence intensity (as Total Radiant Efficiency) in the brain indicated two phases of significant liposome accumulation with an early peak when liposomes were administered around the time of resection surgery (-15’ or +15’) followed by an initial decrease (3h and 24h) and a delayed peak when liposomes were administered 48-72 hours after resection surgery which decreased toward baseline thereafter (**Fig. 1c**). This biphasic liposomal BBB translocation is consistent with that observed in other types of neurological injury such as ischemic stroke^20^. Histological analysis confirmed the results from *ex vivo* whole brain imaging with DiI-liposomes selectively accumulating around the resection cavity/margin (**Fig. 1d**). Notably, while at earlier timepoints (-15’ to 72h) DiI-liposomes were localised around the resection margin, injection on day 7 after resection no longer showed this margin localisation and instead showed high accumulation primarily within residual/recurrent tumour areas that did not extend beyond the tumour borders more consistent with that observed in animals that did not undergo tumour resection (**Fig. S4**). In agreement, the mean area of liposome distribution was significantly reduced by 58% when injected on day 7 compared to the peak spread when injected 48h after resection surgery (**Fig. S5**). This localised accumulation of liposomes was also confirmed in an alternative model based on orthotopic implantation of mouse glioma neurospheres (mGNS) derived from a genetically engineered (sleeping beauty transposon based) GBM model (NRAS;shTP53-GFP;shATRX;luciferase)^26, 27^ (**Fig. S6**). Successful resection of the majority of the tumour mass was confirmed by bioluminescence imaging (BLI) and histology (**Fig. S6a-c**). As with the GL261 model, selective accumulation of DiI-liposomes was observable around the resection site following intravenous administration either 15’ or 48h after resection surgery (**Fig. S6d-e**). Taken together these data confirm the capacity of intravenously administered liposomes to accumulate in the brain at the resection margin of GBM in different preclinical models.

**Fig. 1:**
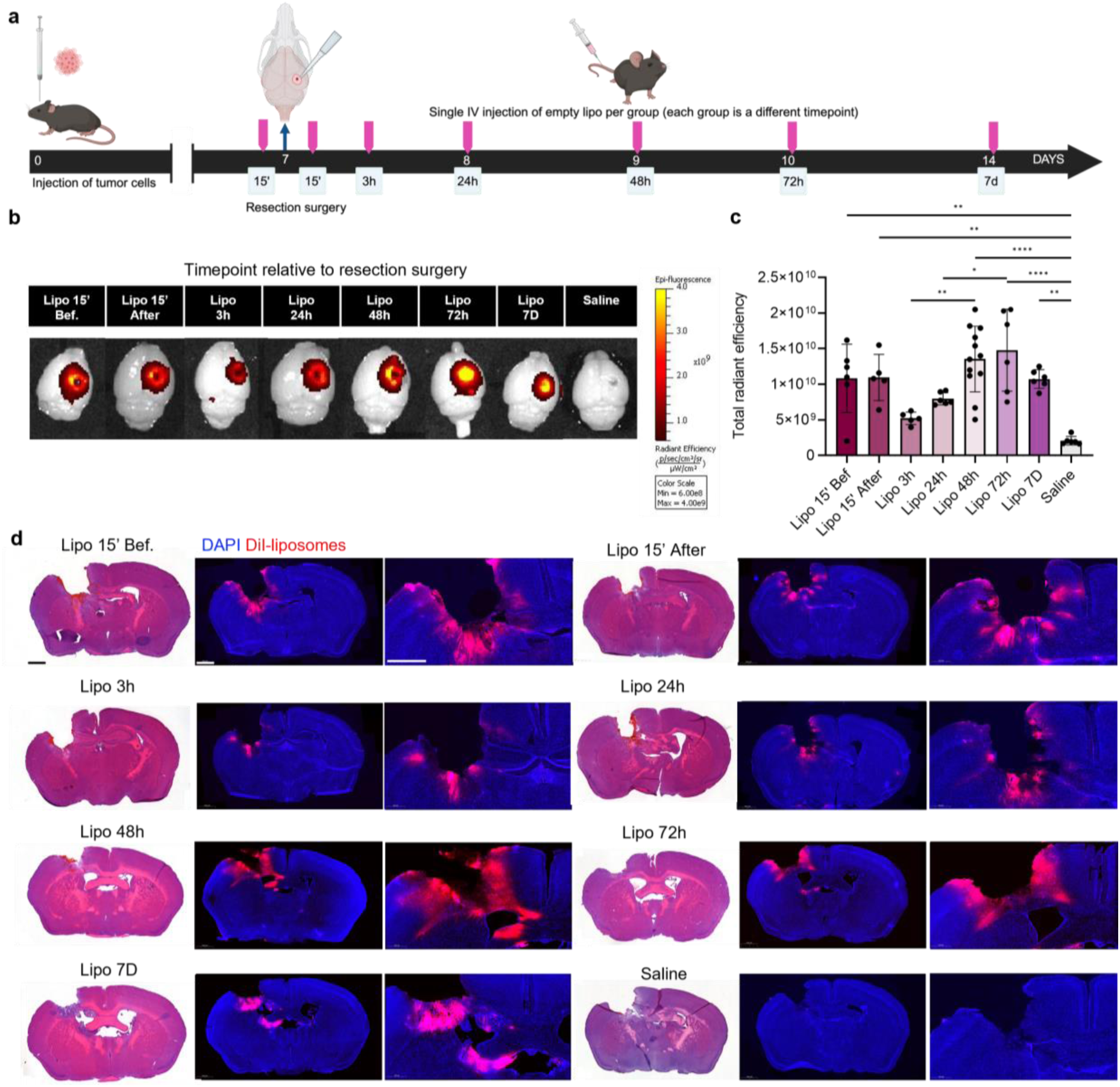
Biphasic accumulation of intravenously administered liposomes in the brain after resection surgery. **a** Schematic of the experimental design and the timepoints of intravenous DiI-liposome injections before or after resection surgery. The time points represent different groups, and each group received a single liposome injection. **b** Representative *ex vivo* images of perfused brains 24h after liposome injection (at the specific timepoints) showing DiI-liposome distribution in the brain around the surgical site. **c** Quantification of liposome fluorescence in the brain expressed as total radiant efficiency (*n* = 6-12). **d** H&E and associated fluorescence microscopy images of brain sections 24h after i.v. injection of DiI-liposomes and selective accumulation around the resection cavity/margin. Scale bars = 1000 µm. Data in (**c**) represents mean ± SD, and *p* values were obtained by One-way ANOVA followed by Tukey’s multiple comparison test. \**p*<0.05, \*\**p*<0.01,\*\*\*\**p*<0.0001

### Postoperative administration of liposomes provides improved uptake into peritumoral/margin areas

To determine the distribution of DiI-liposomes in the postoperative brain we performed spatial analysis of brain sections in animals injected during either the early (15’ after) or delayed (48h after) timepoints of increased liposome accumulation post-resection and compared this to animals that did not undergo tumour resection (**Fig. 2a**). We first delineated the resection cavity (RC) or tumour (T) based on the absence or density of nuclei (DAPI) in resected or unresected control samples respectively (**Fig. 2b**). We then defined the peritumoral area or resection margin (P/M) as the areas within 300 µm of the tumour or cavity edge which equivalates to the area in humans where >80% of recurrence occurs due to the highest presence of residual tumour cells^3, 28^. Animals injected with DiI-liposomes that did not undergo resection showed a high intensity of DiI fluorescence within the tumour area but minimal detectable signal in the peritumoral region (**Fig. 2c**). This is consistent with medical imaging (DCE-MRI) of human GBM which demonstrates a leaky BBB within the tumour bulk (DCE-MRI contrast-enhancing region) with the BBB in the peritumoral regions surrounding the tumour (DCE-MRI non-contrasting enhancing) remaining relatively intact^2, 29^. In contrast, a high intensity DiI-liposome signal could be clearly observed in the P/M region of animals that had undergone resection surgery highlighting local disruption of the P/M BBB and selective accumulation of liposomes at this site. Quantification of fluorescence intensity in the P/M region of animals that were injected with liposomes either 15’ or 48h after resection surgery confirmed a significant increase in liposome accumulation compared unresected animals injected at these same timepoints (**Fig. 2d**). As expected, the DiI fluorescence intensity was significantly lower in the RC compared to the T region of unresected animals, however DiI-Liposomes could be observed to spread beyond the 300 µm margin-zone and into the surrounding brain areas with significantly higher intensity than in unresected animals (**Fig. 2e**). This indicates that postoperative administration of liposomes within these windows of increased accumulation can be used to selectively deliver agents to both margin regions as well as more distant surrounding regions where the more invasive residual tumour cells may reside.

**Fig. 2:**
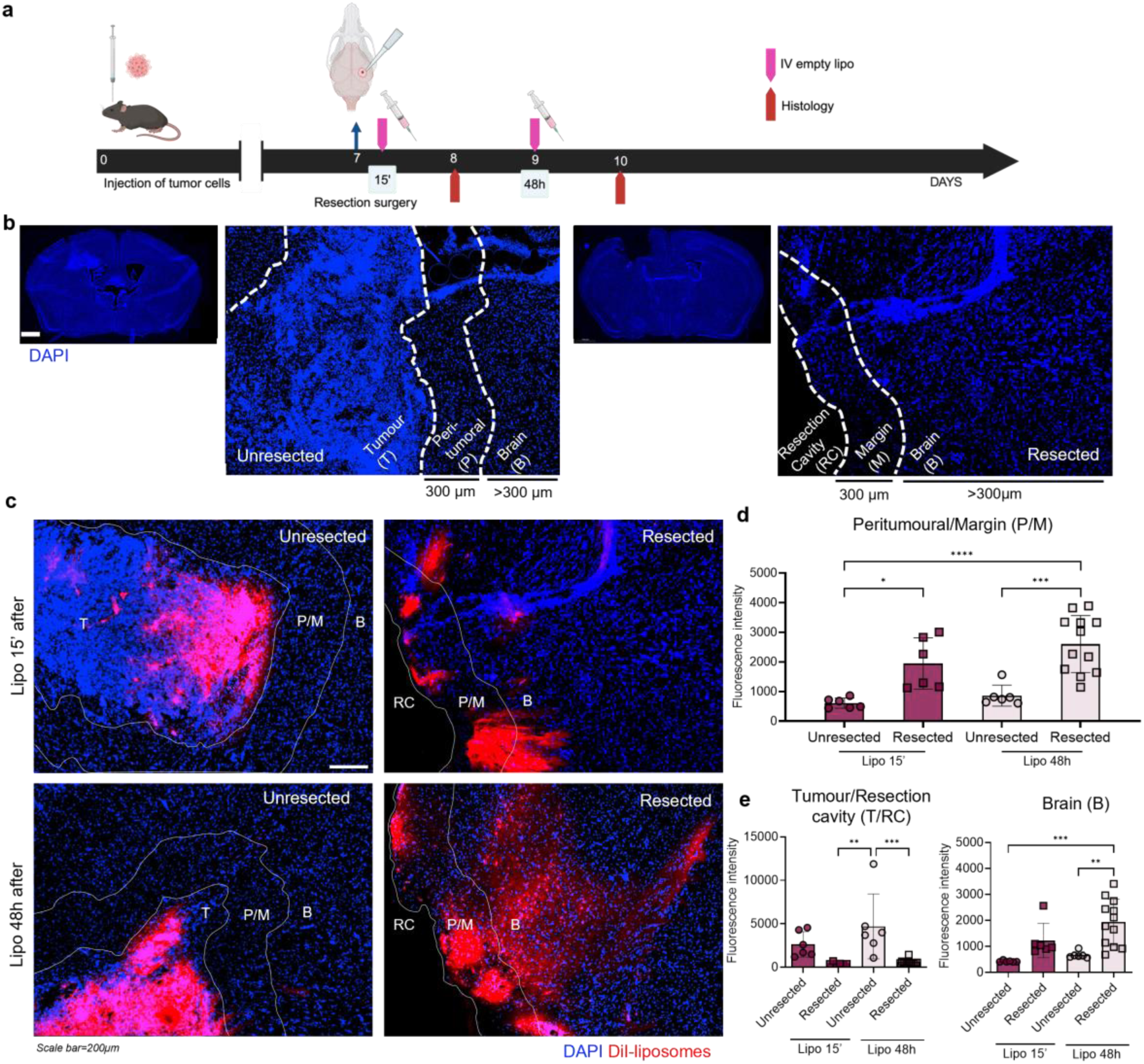
Spatial distribution of intravenously injected liposomes in the resection margin. **a** Experimental schematic. Mice were injected intravenously either 15’ after or 48h after surgical resection of GBM (or at equivalent tumour growth timepoints in unresected animals) before being perfused 24h after injection. **b** Brain areas were delineated based on DAPI appearance and intensity to define tumour (T) or resection cavity (RC) regiones in unresected and resected animals respectively. For both groups, a region of 300µm was considered as either the peritumoral (P) or the resection margin (M). The regions beyond this 300µm were considered as brain (B). **c** Representative images DiI-liposome fluorescence within these 3 regions 24h after intravenous injection at the specified timepoints post-resection (15’ or 48hr) or equivalent in unresected. Scale bar = 200 µm. **d** Quantification of DiI-liposome fluorescence intensity (mean gray value) for the peritumoral/margin region **e** tumour/resection cavity region or brain region (*n* = 6 – 12). Data in (**d,e**) represent mean ± SD (from 3 fields/mouse), and *p* values were obtained by One-way ANOVA followed by the Tukey’s multiple comparison test. \**p*<0.05, **p<0.01, \*\*\**p*< 0.001, \*\*\*\**p*<0.0001

### Spatiotemporal dynamics of BBB disruption to the resection margin can be detected by clinically relevant neuroimaging techniques

We next sought to confirm whether the spatiotemporal patterns of BBB permeability observed using DiI-liposome based optical imaging methods could be detected using dynamic contrast enhanced magnetic resonance imaging (DCE-MRI), a medical imaging technique routinely used in the diagnosis and management of GBM patients. We applied DCE-MRI 24h, 48h and 8 days following resection surgery (**Fig. 3**). Consistent with DCE-MRI in patients with GBM^30^, increased contrast enhancement was present primarily in the tumour area of mice prior to surgical resection, but not beyond the tumour bulk boundary (non-contrast enhancing), indicating a compromised blood-tumour barrier (**Fig. 3a**). In animals that underwent resection, increased signal enhancement extended beyond the resection cavity on days 1 and 2 after resection, with intensity decreasing with distance from the cavity edge (**Fig 3a-c**) indicative of BBB disruption within the resection margin. By day 8, signal enhancement beyond the resection cavity had reduced back to pre-resection levels in agreement with previous data highlighting the accumulation of liposomes only within the tumour area (**Fig 3c** and **Fig 1d**). Overall, these data confirm the presence of resection mediated BBB disruption that, in addition to nanoparticle translocation, can be detected by routinely available MRI techniques.

**Fig. 3:**
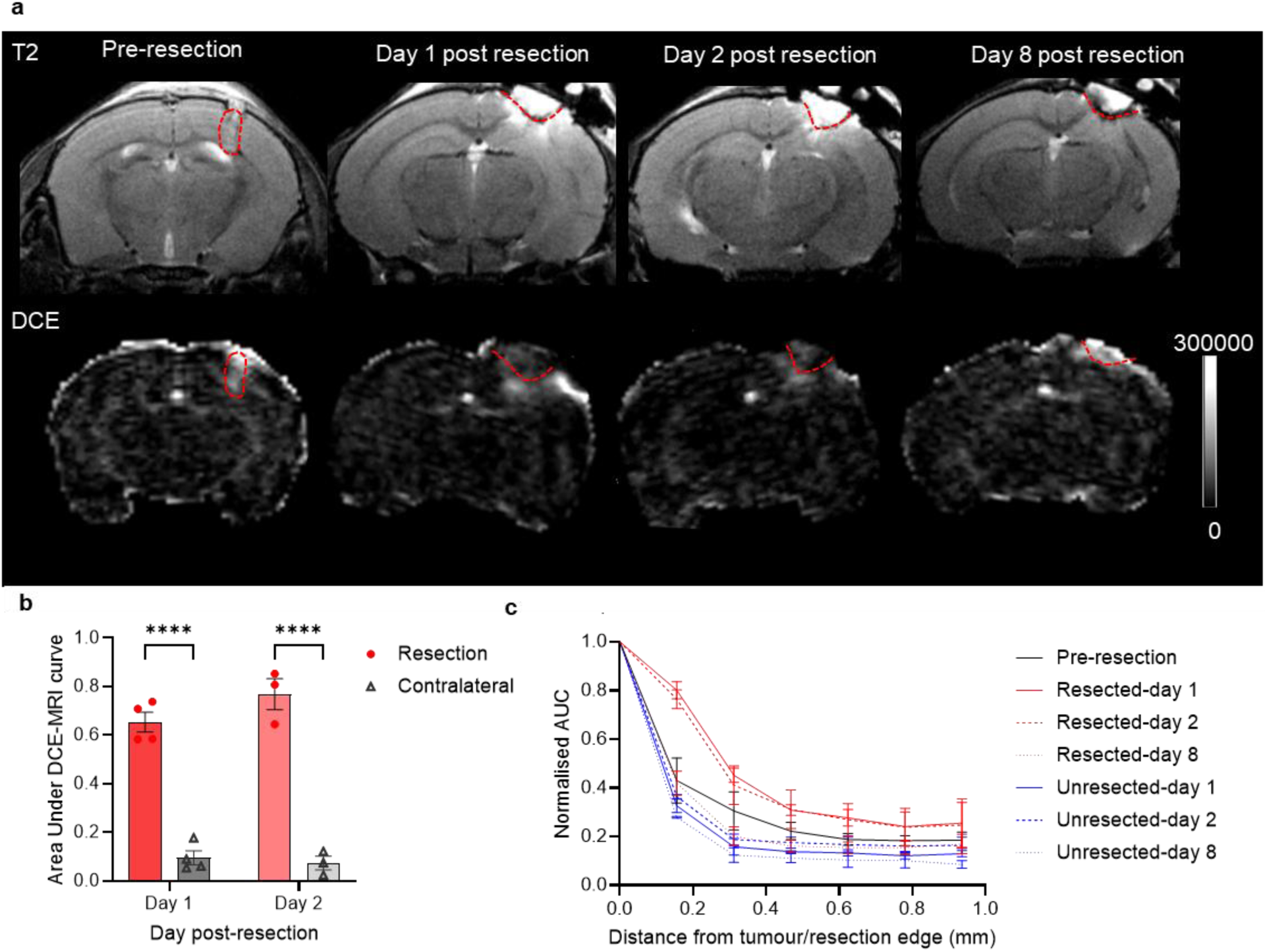
DCE-MRI of resection margin BBB integrity. **a** T2-weighted anatomical and Dynamic Contrast Enhanced (DCE) MRI images of mouse brains before (day -1) and after (24h, 48h, 8D) surgical resection of a GL261 intracranial tumour. Tumour/resection border delineated by red dashed line. **b** Contrast agent leakage, quantified by the area under the DCE-MRI enhancement curve within the resected area and a comparable area in the contralateral hemisphere. **c** AUC, normalised to the AUC within the resection area or tumour, at each 0.2 mm distance from tumour or resection margin edge at each timepoint. Data in **b** and **c** are presented as mean ± SEM. **b** Two-way ANOVA + Sidak-Holm multiple comparisons test. ****p<0.0001

### Translocated liposomes interact primarily with margin localised microglial/macrophage populations

To better characterize the interaction of DiI-Liposomes with the microenvironment of the resection margin, we performed immunostaining of markers associated with blood vessels/endothelial cells (CD31), microglia (IBA1) and astrocytes (GFAP) (**Fig. 4** and **Fig. S7**). We focussed on animals injected with DiI-liposomes on the early (15’) and delayed (48h) timepoints where the most significant accumulation of Dil-liposomes had been observed. 24h after liposome administration and following perfusion of the brains we detected around 40% of DiI-liposomes colocalised with CD31+ blood vessels with the majority of DiI-signal being external to this consistent with extravasation to the surrounding brain parenchyma consistent with ongoing BBB translocation (**Fig. S7a-b**). In agreement with this, DiI-liposomes within the CD31+ cells also showed substantial co-localisation with Cav1, a marker of caveolin-1-dependent transcytosis (**Fig. S7c-d**).

**Fig 4.**
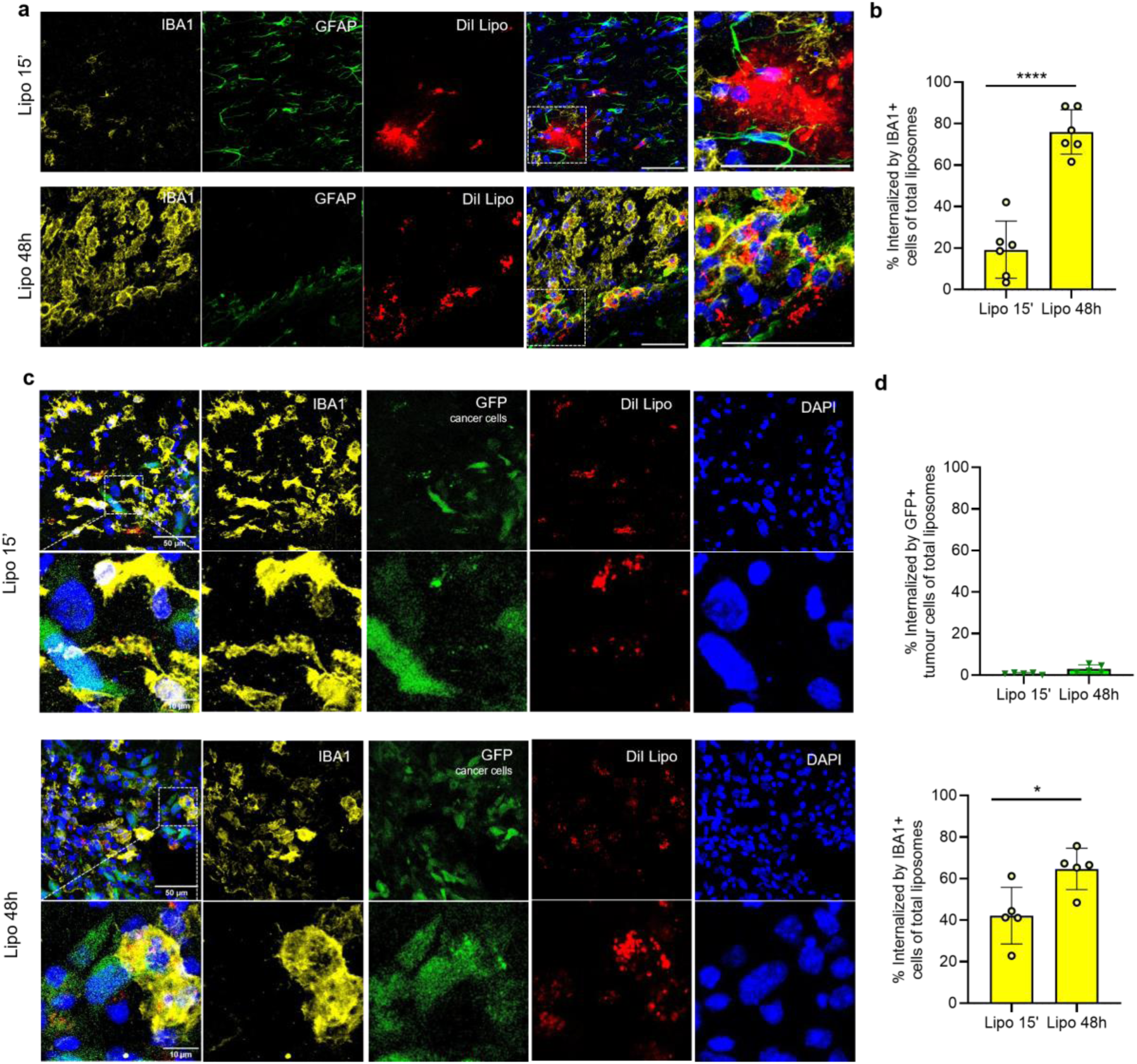
Selective liposome internalization by resection margin macrophages/microglia. **a** Representative confocal micrograph of IBA1 (microglia), GFAP (astrocytes) and DiI-liposomes in mice injected 15’ or 48 hours after GBM resection and brains recovered 24 hours after injection (scale bar = 50 µm). **f** Quantification of internalized DiI-liposomes in IBA1+ cells as a percentage of total liposomes (4 fields of view per mouse, n= 6 mice). **c** Representative confocal micrograph of IBA1 (microglia), GFP (mGNS cancer cells) and DiI-liposomes in mice injected 15’ or 48 hours after GBM resection and brains recovered 24 hours after injection (scale bar = 50 µm; 10 µm). **f** Quantification of internalized DiI-liposomes in GFP+ cancer cells or IBA1+ cells as a percentage of total liposomes (4 fields of view per mouse, n= 5 mice). Data in (**b**,**d**) represent mean ± SD, and *p* values were obtained by Students t-test. \**p*<0.05, \*\*\*\**p*<0.0001.

As we had observed evidence of extravasation we also analysed the interaction of DiI-liposomes with cell types present in the parenchymal microenvironment of the resection margin. DiI-liposomes injected at the early (15’) or delayed (48h) timepoint did not appear to undergo any significant interaction with GFAP+ astrocytes however showed a clear and significant internalisation by IBA1+ macrophages/microglia at both timepoints of injection (**Fig. 4a**). In particular, liposomes injected at 48h post-resection appeared to highly internalised by macrophages/microglia (**Fig. 4b**) which may in part contribute to their distribution and retention in the margin microenvironment at this timepoint. We conducted a similar analysis using the mGNS model, in which glioma cells are labelled with GFP to enable monitoring of their interaction with DiI-liposomes (**Fig. 4c**). Notably, there was minimal internalization of DiI-liposomes by the residual GFP+ cancer cells in the resection margin at either injection timepoint (**Fig. 4d**). Consistent with findings in the GL261 model, the liposomes were predominantly internalized by macrophages and microglia, which showed a close association with the cancer cells within the margin microenvironment (**Fig. 4c-d**). Taken together, these data confirm that intravenously administered liposomes undergo translocation across the BBB within the resection margin and following extravasation interact with resident microglial populations.

### Selective liposome accumulation provides enhanced localised drug delivery to the postoperative resection margin

We next aimed to determine whether this translocation and accumulation of liposomes would translate into an improvement in liposome mediated drug delivery. To assess this, we utilised liposomes encapsulating doxorubicin as the standard and clinically used Doxil^®^/Caelyx^®^ formulation (**Fig. S8**). Doxorubicin, while showing effectiveness on GBM cell lines *in vitro*^31, 32^, has failed to translate due to poor penetration of the BBB and high sensitivity to expulsion by endothelial efflux transporters such that therapeutic concentrations are rarely achieved^33–35^. We administered liposomal doxorubicin (DOX-Lipo or an equivalent dose of free doxorubicin HCl (5 mg/kg) via intravenous administration 48 hours after resection surgery and recovered the brains 1 or 7 days later (**Fig. 5a**). Exploiting the autofluorescent properties of doxorubicin we used confocal microscopy to visualise the distribution of drug in the resection margin. Doxorubicin could be clearly visualised in animals injected with DOX-Lipo but not in animals injected with free DOX (**Fig. 5b**), which was confirmed quantitatively (**Fig. 5c**) and indicated improved drug delivery/retention within the resection margin. We assessed the tissue histologically 24h and 7 days after injection of DOX-Lipo or free doxorubicin and demonstrated that liposome treatment appeared to significantly inhibit the expansion of histologically detectable residual disease compared to free doxorubicin, based on H&E identified tumour volume (**Fig. 5d-e**). Encouraged by this we performed a long-term survival analysis in animals treated with DOX-Lipo or free doxorubicin 48h after resection. Free DOX was unable to significantly delay recurrence compared to vehicle control, whereas DOX-Lipo completely suppressed recurrence up to 100 days in all treated animals (n = 7) in agreement with an enhancement of margin localised drug delivery (**Fig. 5f-h**). These data demonstrated that the selective accumulation of liposomes in the postoperative resection margin can be exploited for enhanced delivery and retention of doxorubicin while offering a rapid treatment strategy to suppress GBM recurrence.

**Fig. 5:**
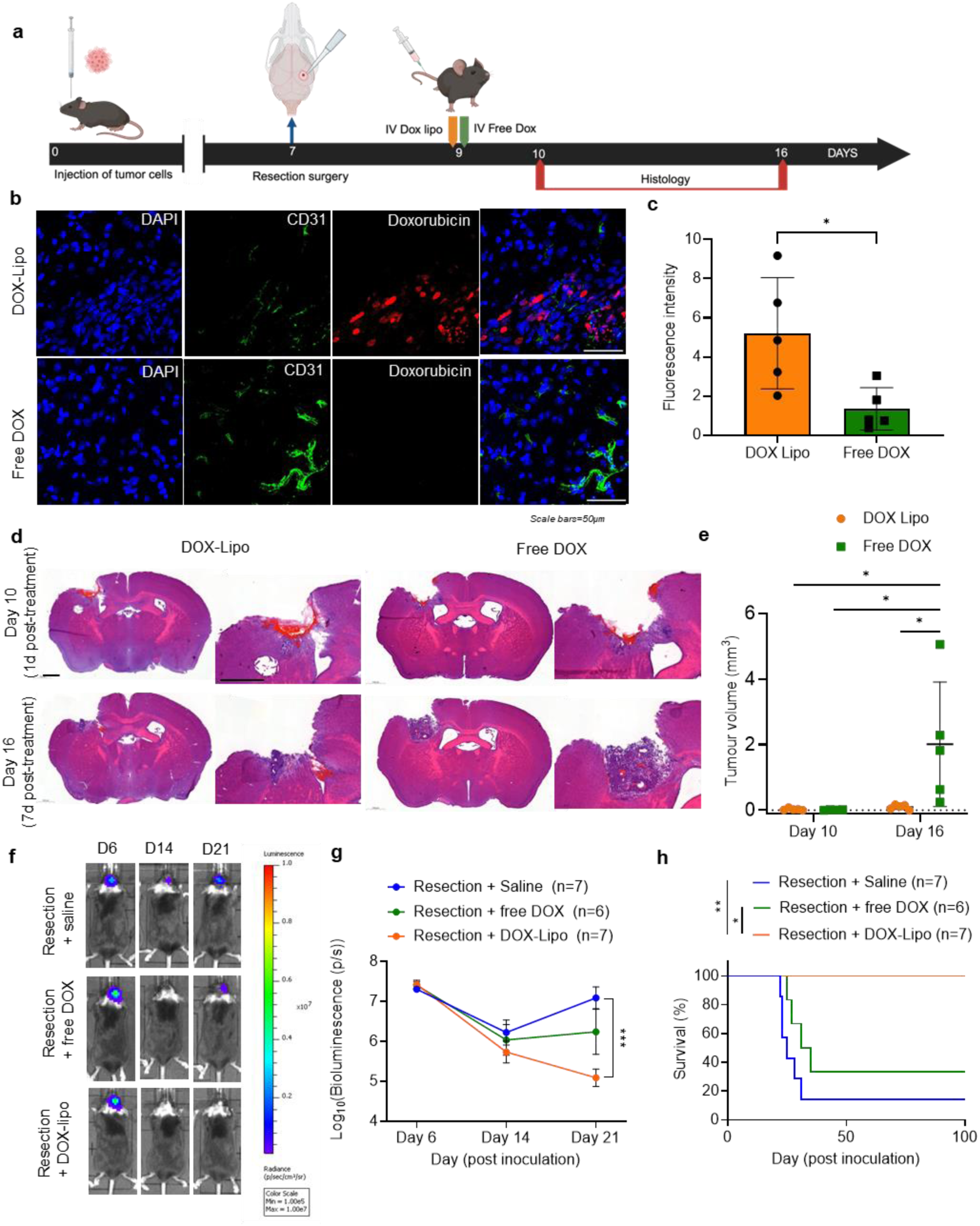
Enhanced resection margin localized liposomal mediated chemotherapy delivery. **a** Experimental schematic. Each group received either liposomal doxorubicin (DOX-Lipo) or free DOX 48h after resection surgery and brains were collected 24h (D10) or 7 days (D16) after treatment. **b** Representative confocal images of doxorubicin autofluorescence and CD31 immunofluorescence in the GBM resection margin 24h after intravenous injection of DOX-Lipo or free DOX injection. **c** Quantification of doxorubicin fluorescence intensity (mean gray value) in each experimental group (3 fields of view per replicate, *n* = 5). **d** Representative H&E histology images of each group with the emergence of early recurrence lesions in free DOX **e** Quantification of residual/recurrent tumor volume in each group (*n* = 5). **f** Representative IVIS bioluminescence images on day 6, 14 and 21 post inoculation (days -1, 7 and 14 post resection) **g** Quantification of tumour bioluminescence (total flux p/s) at each timepoint (n=6-7). **h** Survival curves of animals treated with DOX-Lipo, free DOX or saline control 48h after GBM resection (n=6-7). Data in (**c**,**e**) represent mean ± SD, (g) mean ± SEM. *p* values were obtained by unpaired t-test (c) two-way ANOVA (d, g) followed by Tukey’s multiple comparison test or Log rank (h). \**p*<0.05*, \*\**p*<0.01, \*\*\**p*<0.001, \*\*\*\**p*<0.0001.

We also compared the earlier administration in the immediate postoperative window of BBB permeability (15’) which also achieved a significant improvement in survival over resection alone or DOX-Lipo in the absence of surgical resection (**Fig. S9**). Importantly, early (15’ or 48h) administration of DOX-Lipo treatments did not significantly impact animal health or post-operative recovery in relation to weight or behaviour (**Fig S10**). Overall, these data demonstrate that the early postoperative effects of surgery on the BBB can provide windows for safe and effective application of therapeutic liposomes.

### Early administration of liposomes within the window of BBB permeability is critical to therapeutic outcome

Lastly, to assess the impact of timing of administration and the importance of these identified effects of surgery on BBB permeability to nanoparticles we compared early DOX-Lipo administration (48h) performed above, with a late administration (10 days) that is more typical of the clinical treatment delay with the current accepted SoC (**Fig. 6a**). We observed that while delayed administration of DOX-Lipo (10 days after surgery) did offer some protection against recurrence (4/7 complete responses; CR), this was not significant compared to vehicle control (1/7 CR). In contrast, early administration (48h) offered complete protection from recurrence in all animals (7/7 CR) (**Fig. 6b-d**). Importantly, this underscores the importance of this identified post-operative treatment window for disease outcomes and highlights the potential for the development of margin localised therapeutic strategies to be employed within this time-period.

**Fig. 6:**
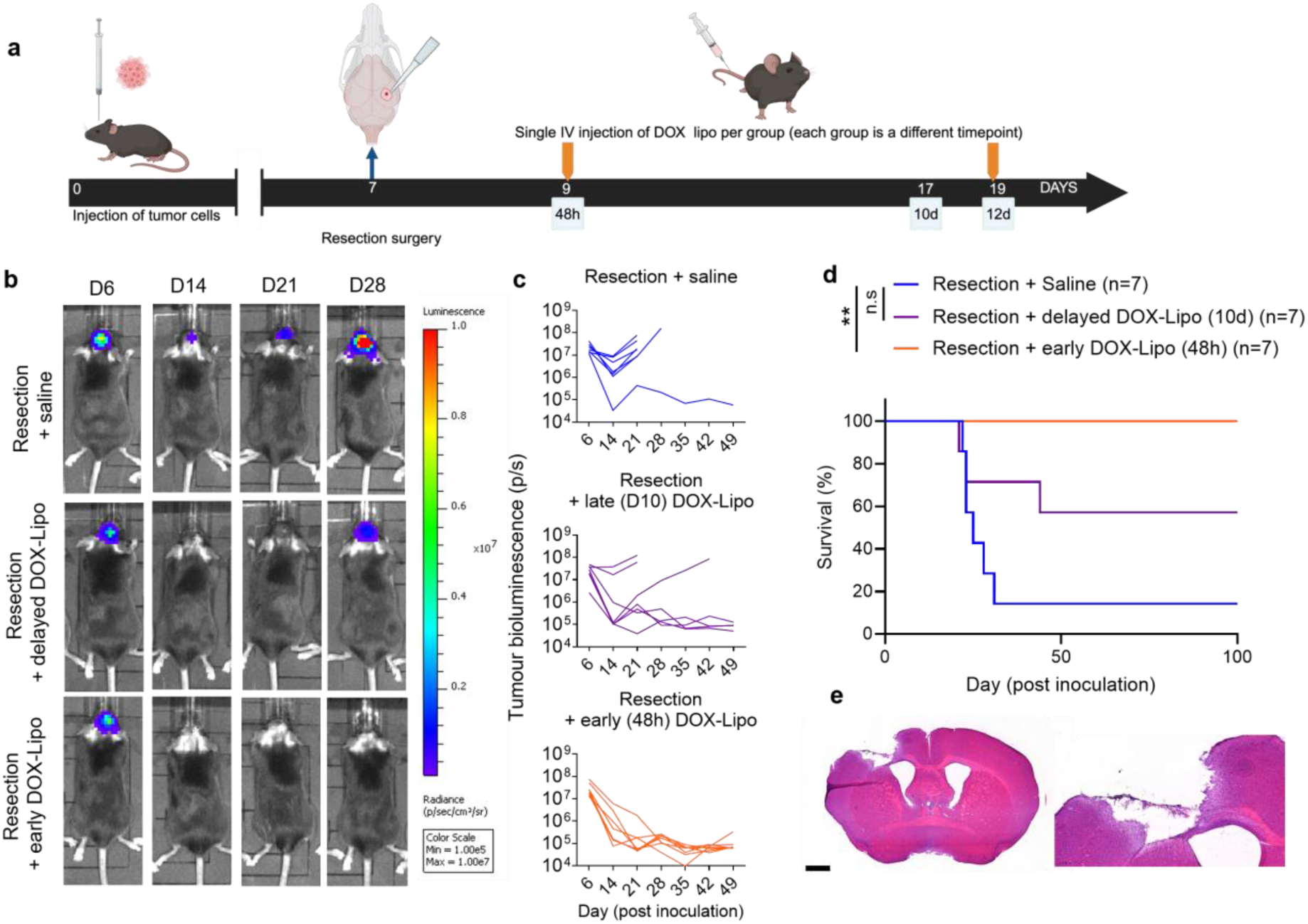
Early administration of liposomes within the window of BBB permeability is critical to therapeutic outcome. **a** Experimental schematic. Following GBM resection mice received intravenous administration of liposomal doxorubicin (DOX-Lipo) either 48h (early) or 10 days (delayed) after surgery. **b** Representative IVIS bioluminescence images on day 6, 14, 21 and 28 post inoculation (days -1, 7, 14 and 21 days post resection) **c** Quantification of tumour bioluminescence (total flux p/s) of individual mice in each group (n=6-7). **d** Survival curves of animals treated with late or early DOX-Lipo or saline vehicle control (n=6-7). **e** representative H&E histology of surviving mouse brain on day 100 (93 days after resection) showing absence of tumour at resection site (n=7). *p* values in **d** were obtained with Log-rank (Mantel-Cox) test \*\**p*<0.01, n.s=not significant.

## Discussion

Effective chemotherapeutic treatment of GBM has consistently been hindered by a functional BBB, limiting the effectiveness of nanoparticle therapeutics, even taking into consideration the higher permeability of the BBB within the tumour bulk. The effects of GBM resection surgery on the blood-brain barrier (BBB) and their implications on chemotherapeutic delivery have not been systematically studied and remain unclear. Our data revealed that surgical resection disrupts the BBB at the peritumoral and resection marginal regions, allowing liposomes to preferentially accumulate in the marginal zone. This dynamic and transient disruption has significant consequences for the effectiveness of post-operative chemotherapy, revealing the opportunity for early treatment initiation within specific temporal windows.

Doxorubicin was initially identified as a promising agent for GBM therapy due to high cytotoxic responses of GBM cells *in vitro*^31^ however, due to poor brain accumulation worsened by doxorubicin clearance^33^, free doxorubicin does not reach cytotoxic levels in brain tumours^36^. Improved doxorubicin delivery has been evidenced with liposomal formulations with clear drug accumulation in unresected tumours mediated by the dysregulated blood-tumour barrier^37^ with similar biodistribution evidence in humans with recurrent GBM^38^. Indeed, several clinical studies have interrogated the potential of liposomal doxorubicin in both newly diagnosed^38–40^ and recurrent GBM^35, 41–45^. Results from such studies have been mixed, although one study showed an improvement in the treatment with liposomal doxorubicin over TMZ alone^40^ and another demonstrated disease stabilization in the recurrence setting^44^. Notably in all cases, liposomal doxorubicin was applied around the time of radiotherapy, at least 4-6 weeks after resection surgery, or in the absence of resection all together. In this way, REP was not avoided which could in part explain the mixed efficacy observed in these trials. In contrast here, by applying liposomal doxorubicin within an identified window of increased BBB permeability to nanoparticles, we demonstrate inhibition of recurrence without any evidence of residual disease in surviving mice.

Different approaches to achieve BBB translocation have previously been explored for post-operative therapy in preclinical GBM. These include transferrin (Tf)-binding, exosome coated nano-micelles, capable of Tf-receptor mediated extravasation into the resection margin^46^. Other strategies have attempted to exploit inflammatory cellular trafficking with drug- or nanoparticle-loaded engineered neutrophils^47, 48^ or platelet membrane-coated heparin-doxorubicin nanoparticles^49^. In the case of engineered neutrophils and neutrophil biomimetics, administration timing proved crucial. Early delivery during the acute post-operative inflammatory response optimized chemotaxis and accumulation at the resection site. Conversely, platelet membrane-coated nanoparticles required pre-operative administration to facilitate optimal infiltration, as post-surgical thrombus formation significantly impeded translocation into the brain parenchyma. Our present study identified two therapeutic temporal windows from surgical resection, an early (within 24hrs post-resection) and a delayed (48-72 hours post-resection) one. Administration of a clinically-used, typically non-BBB penetrant liposome nanoparticle system within such temporal windows, significantly improved therapeutic efficacy. Notably, the use of clinically-established liposomes offers significant translational advantages, including simplified manufacturing processes and an established safety profile in humans, thereby facilitating potential facile clinical translation.

Application of our approach in human patients with GBM will require further exploration of BBB permeability changes in human GBM, and how these changes vary with tumour size and location. In a clinical trial setting non-invasive methods that can measure resection-margin BBB permeability will be necessary to validate optimal timings for liposome administration. We have demonstrated here that routinely available contrast-enhanced MRI can detect and monitor the spatial-temporal dynamics of BBB permeability, indicating the translational potential of these results and highlighting the potential role of DCE-MRI prognostic biomarker for liposomal drug delivery in this setting.

The delay of 4-6 weeks between surgery and the start of chemoradiotherapy has largely been regarded as a necessity to ensure complete recovery from surgery, and is routinely applied in clinical trials of new investigative therapies. However, some clinical studies have demonstrated acceptable safety profiles and favourable outcomes with early (2-3 weeks)^50, 51^ and even ‘super early’ (<7 days)^52^ administration of temozolomide chemotherapy. Furthermore, strategies for immediate intra/postoperative carmustine chemotherapy (Gliadel)^53^ and brachytherapy (Gammatile)^54^ as part of implantable technologies have been clinically approved for GBM and recurrent GBM treatment respectively. Indeed, with the ongoing investigations of neoadjuvant strategies^55, 56^, further exploring how the existing treatment paradigm can be challenged is essential in overcoming the stagnated clinical outcomes in GBM. The early administration of a clinically used liposome formulation within a window of BBB permeability we applied here could be readily implemented within the current SoC and future investigations should determine the safety of this approach in humans.

## Conclusions

Here, we demonstrated that systemic administration of liposomes during newly defined temporal windows of postoperative BBB permeability leads to selective accumulation in the resection margin, a critical target for postoperative GBM therapy. The identification of surgery-induced BBB disruption has significant implications for the design of both preclinical and clinical trials for other poorly BBB-penetrant therapeutic agents and suggests that early administration after resection should be more widely considered. By exploiting this approach for the delivery and accumulation of the clinically used liposomal doxorubicin, we effectively suppressed tumour recurrence in an aggressive resection/recurrence model. We posit that the repurposing of existing, clinically used nanoparticle-based therapeutics can offer a readily translatable approach with the potential to improve the historically poor outcomes in GBM and other resectable brain tumours.

## Experimental

### Empty liposome (DiI-Lipo) synthesis

Liposome vesicles were synthesised from the lipids Hydrogenated soy phosphatidylcholine (HSPC; Lipoid, Germany), Cholesterol (Fisher Scientific, UK), and 2-distearoyl-sn-glycero-3-phosphoethanolamine-N-[methoxy(polyethylene glycol)-2000] (DSPE-PEG2000; Lipoid) at a HSPC:Chol:DSPE-PEG2000 ratio of 56.3:38.2:5.5 mol/mol %. Empty liposomes were prepared by the thin film hydration method followed by vesicle size calibration as previously described^20^. Briefly, lipids were dissolved and mixed in a round-bottom flask in a chloroform/methanol mixture (4:1) with the inclusion of the fluorescent probe 1,1’-Dioctadecyl-3,3,3’,3’-Tetramethylindocarbocyanine Perchlorate (DiI; Invitrogen, 5 mol %). The organic solvents were evaporated to produce a lipid film which was stored at 4 °C overnight, protected from light. Hydration was performed with HBS (20 mM HEPES, 150 mM NaCl, pH 7.4) to a final lipid concentration of 12.5 mM. Extrusion through 800, 200,100 nm polycarbonate extrusion filters (Whatman; VWR, UK) was performed multiple times (5x:5x:20x) to produce unilamellar liposomes with reduced polydispersity. Free DiI was removed by passing liposomes through a Sepharose CL-4B column (Sigma, UK) equilibrated with HBS (pH 7.4) and the final solution concentrated with a Vivaspin column to a final concentration of 12.5 mM liposomes before being sterile filtered through a 0.2 µm membrane.

### Doxorubicin-containing liposomes (DOX-Lipo) synthesis

Doxorubicin was encapsulated in the same liposomal formulation as above (without the fluorescent probe) by following the ammonium sulphate gradient method. The lipid film made of HSPC, CHOL, and DSPE-PEG2000 at 56.3:38.2:5.5 molar ratio, was hydrated in 250mM ammonium sulphate (Sigma) at pH 8.5, at 65°C for 1h. After 1h stabilization at room temperature, the liposomes were extruded with a LIPEX^®^ liposome extruder (Evonik, Germany) at 65°C, five times through 800-400-200nm polycarbonate membranes and five more times through 400-200-100nm membranes. Buffer exchange was performed by Sepharose CL-4B (BioRad, Spain) size exclusion chromatography using HBS and the fractions containing liposomes were further concentrated using Vivaspin 6 (Sartorius) at 8,000g, 20min at 4°C. 0.5 mg mL^-1^ of doxorubicin hydrochloride (Sigma) was added into 12.5mM liposomes (lipid:doxorubicin weight ratio 1:20) and incubated at 65°C for 1.5h. The doxorubicin-containing liposomes were purified using Sepharose CL-4B columns and further concentrated using Vivaspin columns to obtain 12.5 mM liposomes. The final solution was sterile filtered through a 0.2 µm membrane. Doxorubicin encapsulation efficiency resulting of 98% was obtained using 0.1% Triton X-100 (TX) to dissolve the liposomes and the doxorubicin fluorescence was quantified using the plate reader SpectraMax® iD3 at ICN2 Nanobioelectronics and Biosensors Group, excitation and emission wavelengths of 480 and 593nm, respectively, and calculated as:

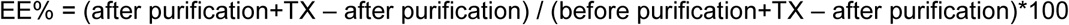

### Cell lines and culture conditions

GL261-luc cells were kindly provided by Prof Brian Bigger (University of Manchester). GL261 cells were expanded in RPMI (Gibco) supplemented with 10% fetal bovine serum (FBS, Sigma Aldrich) and 1% penicillin and streptomycin (P/S) (Gibco). The murine glioma neurosphere line (mGNS; NRAS;shTP53-GFP;shATRX;luciferase) was kindly provided by Prof Maria Castro, University of Michigan. These cells were cultured as undifferentiated neurospheres in DMEM/F12 (Gibco) supplemented with 1X B-27 (Gibco), 1X N2 (Gibco), 1% P/S, 20 ng/ml mEGF (Peprotech, UK) and 20 ng/ml mFGF (Peprotech, UK). Cells were maintained at 37 °C, 5% CO_2_ in a humidified environment. For inoculation, cells were detached/dissociated using Accutase (Thermo Scientific) and washed 2x in PBS to remove media components.

### Animals

All animal experiments were performed at the University of Manchester (UK), in accordance with the Animals (Scientific Procedures) Act 1986 (UK), approved by the University of Manchester Ethical Review Committee and under a UK Home Office Project License P089E2E0A/PP3096857. Animals were housed in groups within ventilated cages with *ad libitum* access to food and water. C57BL/6 mice were purchased from Charles River, UK and were allowed to acclimatize to the facility for at least one week prior to any procedure.

### Intracranial inoculation of glioma cells

Mice were inoculated with syngeneic GBM cells as previously described^57^. Adult female C57BL/6 mice (10-11 weeks old) were anesthetized using isoflurane (2.5% induction and 1–2% maintenance in medical oxygen, at a rate of 1.5 L min−1) and placed on a stereotactic frame. Prior to incision, animals received 0.1 mg kg^−1^ of buprenorphine (Buprenex, Reckitt Benckiser, UK). A midline incision was performed to expose the cranium and a 0.7 mm borehole was drilled (Fine Science Tools, Canada) above the right striatum at 0.7 mm anterior and 2.4 mm lateral from bregma. A 10 μl Hamilton syringe (SYR10, Hamilton, USA) fitted with a 26-gauge blunt needle (Hamilton, USA) was lowered to 1.5 mm below the cortical surface and slowly withdrawn 0.5 mm such that the injection took place at 1.0 mm depth. 5 × 10^4^ GL261-luc or mGNS cells in 1 μl of PBS were injected slowly over 5 min at a rate of 0.2 μl min^−1^. Post-injection the needle was kept in place for 3 min to minimize reflux and slowly withdrawn over 1 min to minimize any injury. The skin incision was closed with 6-0 coated vicryl sutures (Ethicon, UK) and animals were allowed to recover in a heated environment.

### Resection surgery

Tumour-bearing mice were anesthetized using isoflurane (2.5% induction and 1–2% maintenance in medical oxygen, at a rate of 1.5 L min^−1^) and placed on a stereotactic frame. Prior to incision, animals received 0.1 mg kg^−1^ of buprenorphine (Buprenex, Reckitt Benckiser, UK) and 0.1 mg Kg^-1^ dexamethasone. A midline incision was performed to expose the cranium near the previous borehole made. A craniotomy was performed using a dental drill (NSK-Nakanishi, Japan) to expose the brain at the tumour site. The dura was incised and the tumour was resected via aspiration using a vacuum pump (Integra VACUSIP, VWR) and resection of all visible tumour mass was performed in all experimental groups. The craniotomy skull cap was placed back above the brain tissue and sealed with Kwik-Sil (WPI, USA). Finally, mice were sutured, provided with 0.9% saline, and allowed to recover in a heat-box with mash food and continuous monitoring.

### Liposome Biodistribution

Tumour bearing mice were intravenously injected with 200 mL of DiI labelled liposomes in HEPES at a single timepoint either -15’ before resection, 15’, 3h, 24h, 48h, 72h or 7 days after resection, or sham surgery. 24h after liposome injections, mice were anesthetized with 2.5% isoflurane and culled by cardiac perfusion with 2 mM EDTA in PBS to completely remove blood and circulating liposomes. Their brain and organs (heart, lungs, liver, kidneys and spleen) were removed and placed on Petri dishes for IVIS fluorescence imaging (IVIS Lumina II, PerkinElmer, UK) of the liposomes 549 nm excitation, 565 nm emission. Images were analyzed with Living Image software (version 4.7) (PerkinElmer, UK). Following imaging, brains were collected and fixed in 4% PFA for further histological analysis.

### Doxorubicin distribution

GL261-luc bearing mice were treated with 200 mL of Doxorubicin-containing liposomes (5 mg kg^-1^) or free doxorubicin (5 mg kg^-1^) IV at a single dose 15’ or 48h after resection surgery. 24h or 7 days after liposome treatments mice were anesthetized with 2.5% isoflurane and culled by cardiac perfusion with 2mM EDTA in PBS. Following tissue sectioning, doxorubicin was visualized using its intrinsic fluorescence (excitation 470 nm; emission 595 nm), with a Leica SP8 inverted WLL confocal microscope. Fluorescent doxorubicin was measured by mean grey value on single doxorubicin channel from Z-stack 2D images. The threshold of greyscale image was applied to subtract non-specific background.

### Therapeutic treatment design

Doxorubicin-containing liposomes (5 mg kg^-1^), free doxorubicin (5 mg kg^-1^) or vehicle control were intravenously injected at a single timepoint either 15 minutes, 48h or 12 days after resection (or sham) surgery. IVIS, monitoring and histological analysis were performed in a blinded manner. Animals were monitored and weighed daily and tumour re-growth was evaluated by IVIS bioluminescence imaging (BLI) weekly. Animals were sacrificed when they reached humane-end points related to weight loss or the onset of neurological symptoms and brains were collected for histology. Initial BLI measurements were used to group animals randomly such that each treatment group had equal mean starting tumour size. BLI and animal monitoring was performed blinded.

### In Vivo Bioluminescence Imaging (BLI)

Tumour-bearing or tumour resected mice underwent intraperitoneal injection with 150 mg kg^−1^ mouse D-luciferin (15 mg ml^−1^; Promega, UK) in PBS followed by anaesthesia with 1.5% isoflurane. After 8 min, bioluminescence signals were detected using sequential imaging (15 measurements at 2 min intervals) with an in vivo imaging system (IVIS Lumina II, PerkinElmer, UK). Images were analysed with Living Image software (version 4.7) (PerkinElmer, UK). Tumour bioluminescence (photons/s) was log-transformed and statistical analysis performed.

### MRI

Tumour bearing mice underwent MRI on a 7.0T Agilent magnet interfaced to Bruker Avance III console. Mice were scanned at baseline, 6 days after tumour inoculation. On day 7, n = 4 tumour-bearing mice were submitted to a resection surgery and assessed with MRI 24h, 48h and 8 days following surgery. n=2 mice from the initial group did not undergo resection and were scanned with MRI at the same timepoints as resected mice to assess BBB permeability in non-resected mice. Each MRI session consisted of T2-TurboRARE MRI and DCE-MRI, Quantities, processes and model definitions used to report DCE-MRI acquisition and analysis are OSIPI CAPLEX compliant^58^. T2-TurboRARE MRI (TR/TE = 3000/35 ms, FOV = 20 x 20 x 7.5 mm, matrix size = 256 x 256 x 20) was performed to define the anatomical position of the tumour or resection margin. Dynamic contrast-enhanced MRI using 3D spoiled gradient echo (TR/TE = 8.0/1.6ms, flip angle = 12°, FOV = 20 x 20 x 7.5 mm, matrix size = 128 x 128 x 10, temporal resolution = 10.2s, duration = 20 min 28s) was used to measure BBB integrity, by monitoring the increase in DCE-MRI signal intensity caused by leakage of contrast agent from blood into brain tissue. Each mouse was anesthetised using 4% isoflurane in 100% O_2_, cannulated using a 27G needle attached to a line containing Gd-DOTA contrast agent, and positioned on the MRI bed. Once stable, anaesthesia was reduced to 1.5-2%. Ear and bite bars were used to secure the head. A volume resonator was used for transmission and mouse brain surface coil for reception. After the 18^th^ dynamic frame (3-minute baseline), 0.1mmol/kg of Gd-DOTA was injected as a bolus using a contrast injector at 1 mL min^-1^. Following completion of MRI, mice were recovered in a heated box and returned to housing.

### MRI Analysis

Tumour and resection margin regions of interest (ROIs) were delineated manually on T2-TurboRARE images in MRIcron (version 1.0.2). These ROIs were imported into Matlab (Mathworks, version 2023a) where they were down-sampled to the grid size of the DCE-MRI data. Prior to applying to the DCE-MRI data, a map of DCE-MRI area under the curve (AUC) was generated from each dataset. First, DCE-MRI timeseries were coaligned to remove any motion between frames. Second, the DCE-MRI signal was converted to signal enhancement by subtracting the baseline signal. Third, the signal was summed across all timepoints to calculate the AUC. Once the AUC maps were calculated, the tumour or resection ROI was applied to extract the mean AUC. To calculate the AUC at different distances from the resection or tumour margin, the original ROI was dilated by 1 pixel in all direction, then subtracted from the original ROI leaving a layer of ROI pixels just outside the original ROI. Any part of this layer residing outside of the brain tissue was excluded. The mean AUC for this layer was then extracted, and the process repeated a further 5 times, providing a profile of AUC outwards 1mm from the resection or tumour edge.

### Tissue Processing and Staining

At the end of each experiment, mice were anesthetized with 2.5% isoflurane and culled by cardiac perfusion with 2 mM EDTA in PBS to completely exsanguinate blood and circulating liposomes. Brains were removed and fixed overnight at 4 °C and later placed in 30% sucrose in PBS for at least 24 h. The brains were snap-frozen in cold isopentane (−40 °C) and coronal sections (20 μm thickness) were taken using a cryostat (Leica CM1950, Leica Biosystems, Germany).

### Histological Evaluation of Tumor Growth

Sections were stained with haematoxylin and eosin (H&E) staining to observe the histological characteristics of the tumor sections and determine the tumor volume. Cryosections were left for 15 min to air dry before fixation with pure ethanol for 2 min. Slides were washed once with PBS for 5 min, and were placed in haematoxylin for 1 min. Slides were washed twice with water for 3 min each and were placed in 70% EtOH for 3 min. Following dehydration, slides were placed in eosin solution (1% eosin in 95% alcohol) for approximately 40 s. This was followed by three washes in 100% alcohol for 3 min each, and slides were placed in two changes of Xylene for 2 min each. Finally, DPX mount was used to mount coverslips and slides were then left to dry overnight at room temperature. Slides were scanned using a 3D Histech Panoramic 250 slide scanner. H&E stained sections were imaged using a Panoramic 250 slide scanner (3D Histech, Hungary) and analyzed using 3DHISTECH Case Viewer software version 2.6. Initially, tumor diameter was measured in each section so as to identify the maximal tumor area. Subsequently, the height and width of the tumor area were measured and the volume was calculated using the following formula: V = (W^2^ × H) ∕2.

### Immunofluorescence (IF) Staining and Analysis

For immunofluorescence analysis, 20 μm cryo-sections samples were air-dried and fixed for 10 min in ice-cold acetone. After washing the samples with PBS, sections were incubated for 1 h with 1% bovine serum albumin and 5% goat serum in PBS-Triton X 0.2% to remove any non-specific binding. Rabbit anti-IBA1 antibody (dilution 1:500, Fujifilm, 019–19741 Wako), chicken anti-GFAP antibody (dilution 1:1000, Millipore, AB5541), rat anti-CD31 antibody (dilution 1:100, BD Biosciences, 550274) and mouse anti-Cav1 antibody (dilution 1:100, BD Biosciences, 1, 610407) were incubated overnight at 4 °C. For secondary antibody staining, Alexa Fluor™ 647-conjugated goat anti-rabbit IgG (dilution 1:400, Invitrogen, A21244), Alexa Fluor™ 488-conjugated goat anti-chicken IgY (dilution 1:500, Invitrogen, A11039), Alexa Fluor™ 647-conjugated goat anti-rat IgG (dilution 1:200 Invitrogen, A-21247) and Alexa Fluor™ 488-conjugated goat anti-mouse IgG (dilution 1:200, Invitrogen, A-11001) were used. DAPI (Merck, UK) was also added to sections in a 1:10000 dilution for nuclei staining. Sections were washed and Prolong Gold medium (Thermo Scientific) was added and covered with coverslips. Images were taken with a Histec Pannoramic250 slide scanner, epifluorescence (Zeiss AxioImager Z1 fluorescence) and confocal (Leica SP8 WLL inverted confocal) microscopes. Images were processed and analysed using ImageJ for measurements of fluorescence intensity, co-localisation and internalisation.

For co-localisation analysis maximum intensity Z-projection was used to reconstruct blood vessels in 2D from confocal images strained with CD31 and/or Cav1. Dual-color images from two selected channels were converted to RGB format, and colocalization was assessed using a color threshold method. The areas where the two colors overlapped were identified as colocalized regions, with the extent of colocalization expressed as a percentage.

For internalization analysis (DiI-IBA1, DiI-GFAP, DiI-GFP) cell populations were manually counted via cell counter plugin from using 3D multiple-focal plane images and internalized DiI-liposomes were quantified per field of view. Data was expressed as internalized DiI-liposomes within cell population as a percentage of total DiI-liposome signal present across all populations.

### Statistical Analysis

Statistical analysis was performed using GraphPad Prism 10. For the comparison of two groups, an unpaired Student’s t-test was used and for comparison of three or more groups, an ordinary one-way ANOVA (Tukey’s multiple comparison test) or two-way ANOVA (Tukey’s or Sidak’s multiple comparisons test) were utilized. For survival curves, Log-rank (Mantel-Cox) test was performed. Data was regarded as statistically significant if p < 0.05. p-values and statistical tests are specified in the figure legend for each data. Data are presented as mean ± SD or S.E.M as defined in the figure legend of each figure.

## Author Contributions

L.F. T.K performed *in vivo* experiments, developed methodology, performed data analysis, figure preparation and wrote the manuscript. C.P. performed tissue staining and imaging, performed data analysis and figure preparation. B.D. contributed to methodology development and experimental design of MR imaging, performed data analysis and provided input to interpretation of data and figure preparation. Y.H. and L.T. assisted with experiments, *in vivo and ex vivo* imaging, and tissue processing. L.F. and N.L. contributed to the preparation and characterization of the liposome nanoparticles. N.H performed liposome cryo-EM sample preparation and imaging. L.F., K.K. and T.K. conceptualized and designed the study. K.K. and T.K. coordinated the study and acquired funding. L.F., B.D., N.L., K.K and T.K. contributed to the writing of the manuscript.

## Acknowledgements.

The authors would like to acknowledge Mang Xu, Alexandra Thawley and Kiana Arashvand for assistance with tissue processing and histological preparations. The authors would also like to thank the Histology core facility, Bioimaging core facility, Electron microscopy facility and the Biological Services facility at the University of Manchester for technical support and access to equipment. NL would like to acknowledge the ICN2 Nanobioelectronics and Biosensors Group (led by Prof. Arben Merkoçi) for the access to the plate reader. NH would like to thank Mr. Martí del Cabo from the Microscopy and X-Ray Diffraction Service at UAB for assistance on the sample preparation and microscope data acquisition. TK would like to acknowledge the University of Manchester/Medical Research Council (MRC) Confidence in Concept Round 8 for financial support. TK and KK would also like to acknowledge the Engineering and Physical Sciences Research Council (EPSRC) 2D-Health Programme Grant (EP/P00119X/1) and EPSRC International Centre-to-Centre grant (EP/S030719/1). ICN2 is funded by the CERCA programme / Generalitat de Catalunya and is supported by the Severo Ochoa Centres of Excellence programme, Grant CEX2021-001214-S, funded by MCIU/AEI/10.13039.501100011033. The authors would also like to acknowledge Professor Maria Castro (University of Michigan) for provision of the mGNS cell line that was derived with support from NIH/NINDS 1RO1NS122165-O1.

