## Supporting Information for "Targeting therapeutic nanoparticles to the glioblastoma resection margin by harnessing post-operative spatiotemporal blood-brain barrier disruption"

---

* Correspondence to be addressed to either:


### Supplementary Experimental

**Liposome characterization.** Liposome systems fabricated were characterised using the following techniques:

**Dynamic light scattering and zeta potential measurements**. Liposome size (hydrodynamic diameter) and surface charge (zeta potential) were measured using a Zetasizer Nano ZS (Malvern, Instruments, UK). Samples were diluted 100 times with distilled water and 3 independent measurements per sample were recorded, and the data expressed as average ± SD.

**Transmission electron microscopy (TEM).** Empty DiI-labelled liposomes were visualized with transmission electron microscopy (FEI Tecnai 12 BioTwin) at a 1 mM lipid concentration with a Carbon Film Mesh Copper Grid (CF400-Cu, Agar Scientific). Samples were stained with 1% aqueous uranyl acetate solution.

**Cryo-transmission electron microscopy (Cryo-TEM)**. Doxorubicin-containing liposomes were imaged with a Cryo-transmission electron microscopy (CryoTEM) at the Microscopy and X-ray diffraction service at UAB, using a JEOL-2011 transmission electron microscope adapted for cryogenic microscopy, working at a voltage of 200kV and equipped with CMOS Gatan Rio 16 camera and EDS Oxford Instruments X-max detector.

### Supplementary Figures

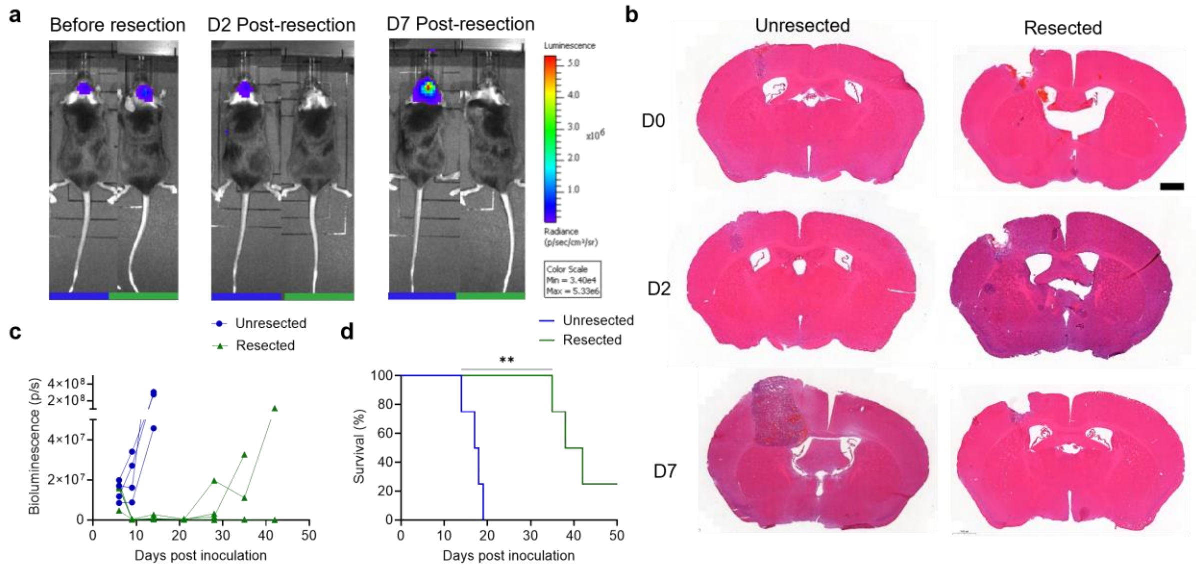


**Supplementary Fig. 1: Establishment and characterization of a GBM resection/recurrence model. a** Longitudinal *in vivo* imaging of tumor bioluminescence 1 day before (day 6) and 2 or 7 days after (day 9 and 14) resection surgery. **b** H&E histology images of unresected and resected brains on days 0, 2 and 7 after resection (days 7, 9 and 14 of tumour growth). Representative images from n=4. Scale bar = 1000 µm. **c** Quantification of total tumor bioluminescence from each animal of the unresected and resected groups overtime (*n* = 4). **d** Survival (humane endpoint) curves of mice with or without tumour resection (n=4). Log-rank test. **p<0.01

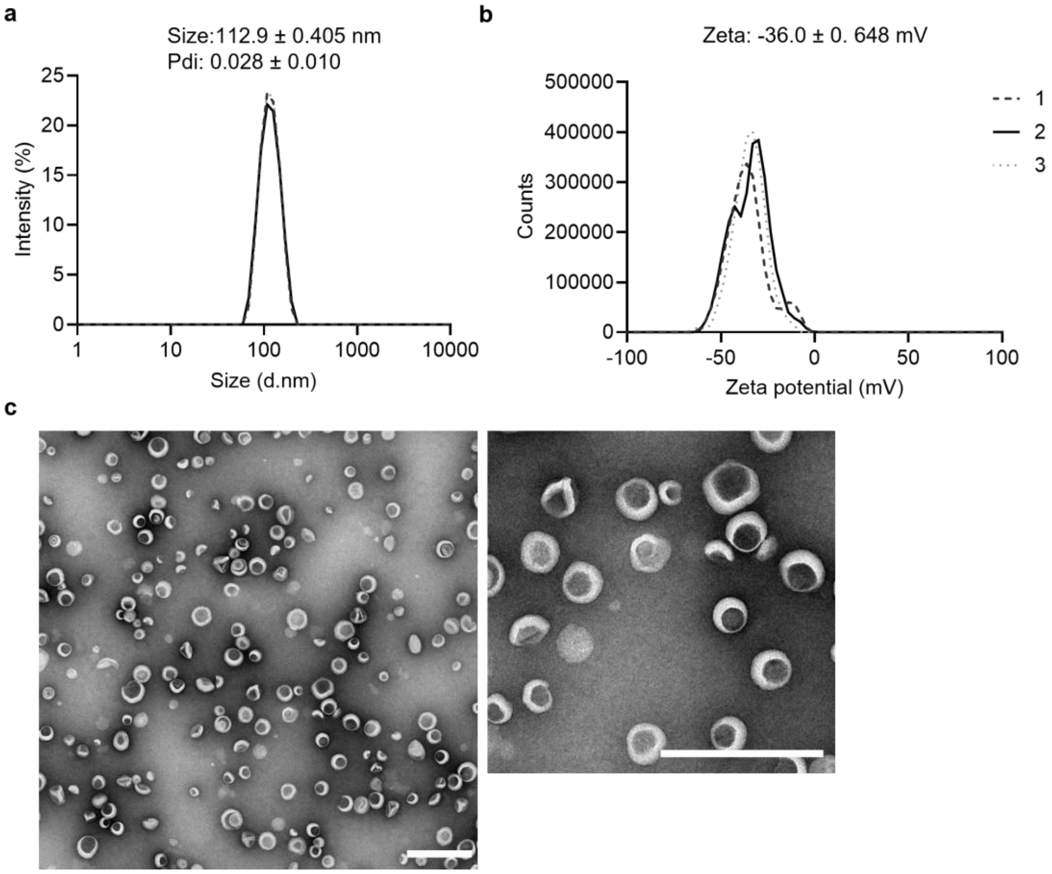


**Supplementary Fig. 2: Characterization of empty DiI-labelled liposomes. a** Dynamic light scattering (DLS) measurement of size distribution and **b** zeta potential of empty-DiI labelled liposomes. Data is represented as the mean ± standard deviation of 3 measurements. c empty-DiI labelled liposomes with TEM. Scale bar = 500 µm.

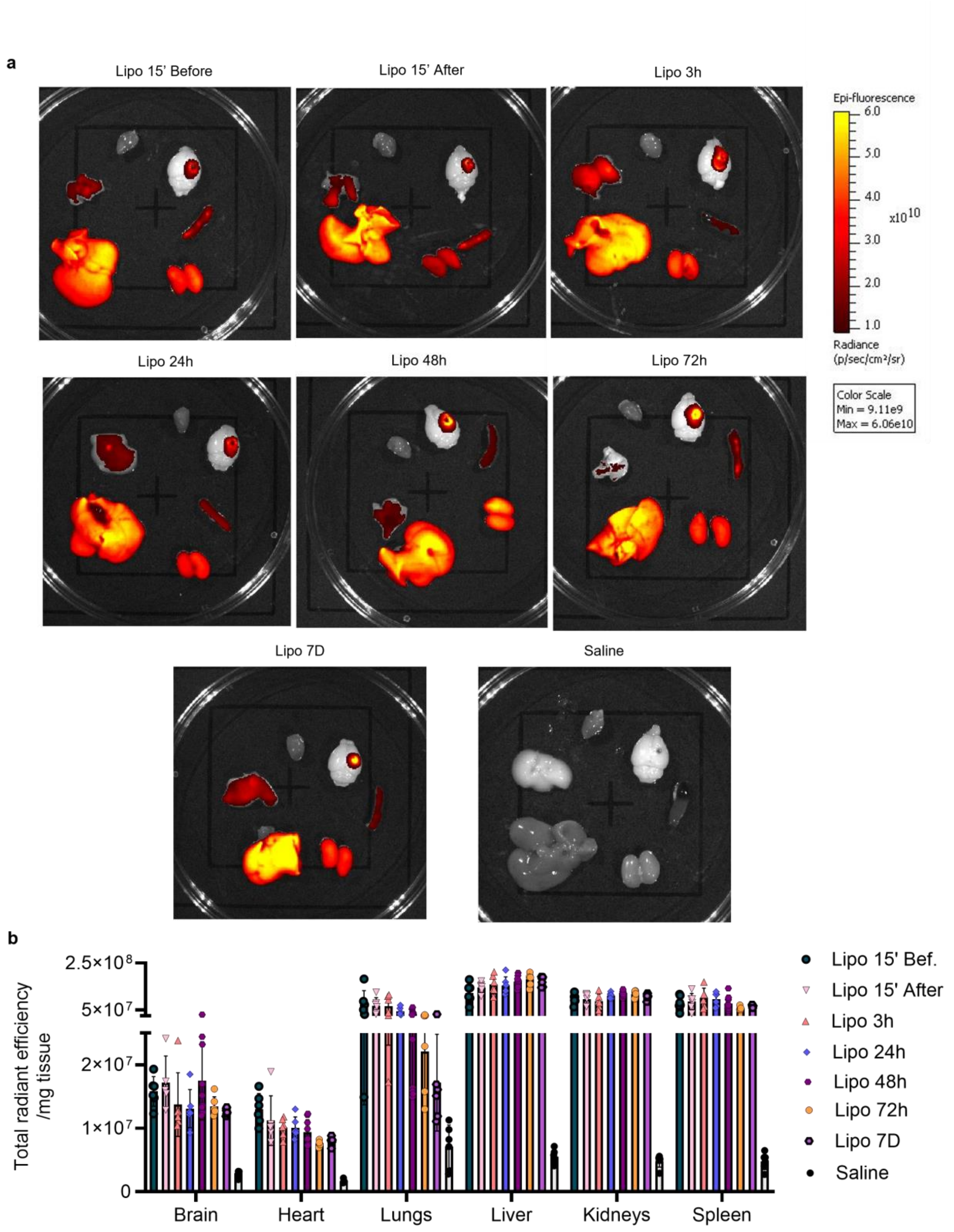


**Supplementary Fig. 3: Biodistribution of intravenous empty DiI-liposomes injected at different timepoints. a** Ex vivo imaging showing liposome fluorescence within organs of interest obtained using IVIS Lumina II imaging system at each of the injection timepoints (24h after injection at the specified timepoints). **b** Quantification of liposome DiI-fluorescence per mg of tissue in each organ/injection timepoint (n=6-12).

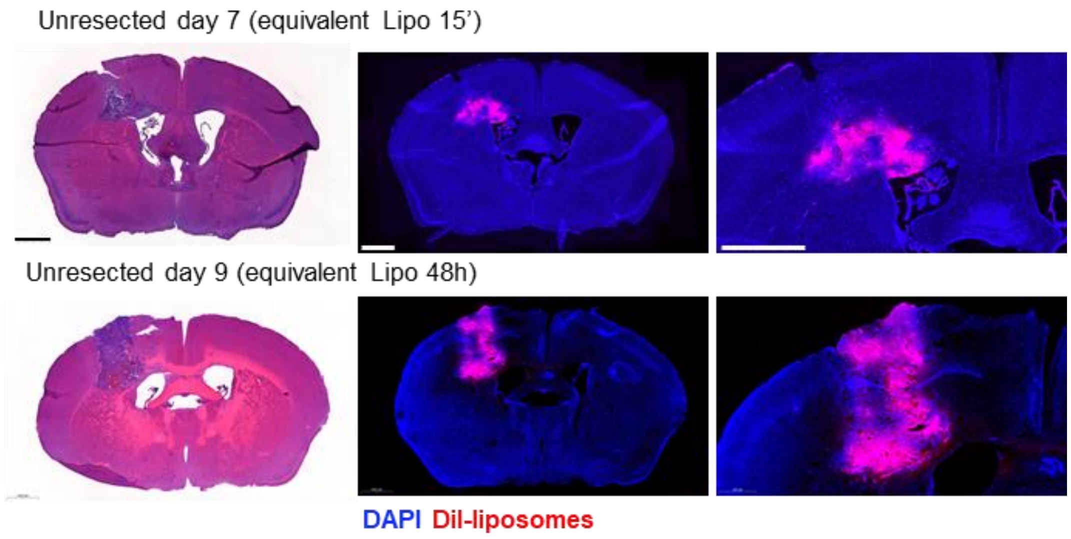


**Supplementary Fig. 4: Tumour accumulation of empty DiI-liposomes in the brain of unresected groups.** Representative H&E and fluorescent microscopy images showing the distribution of DiI-liposomes on perfused brain sections of GL261 tumour bearing animals following injection at the specific timepoints. Brains were collected 24h after i.v injection. Scale bars = 1000 µm.

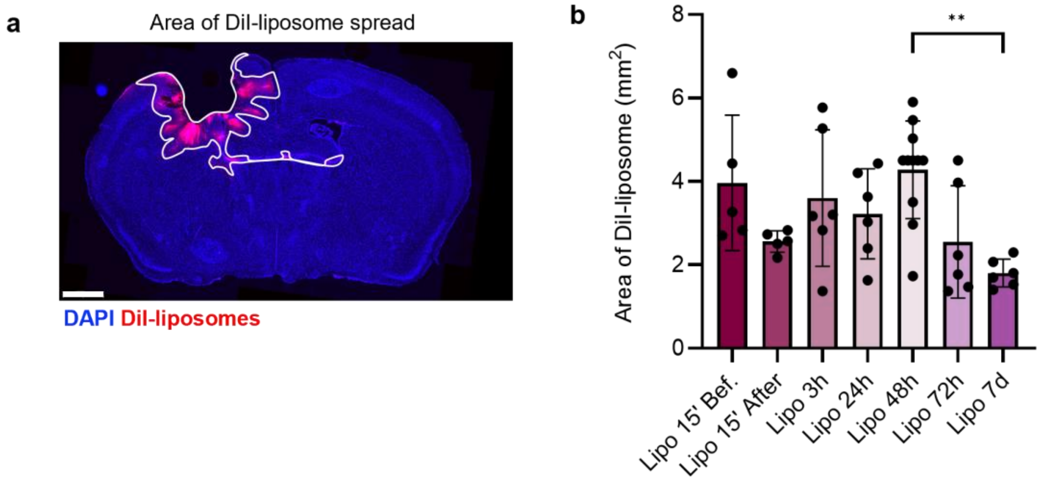


**Supplementary Fig. 5: Quantification of the area and distance of DiI-liposomes distribution around the resection margin. a** Schematic representation of the liposome distribution area analysis performed on brain sections from animals administered with liposomes -15'. +15'. 3h 24h 48h, 72h or 7 days after resection surgery (images from 24h after injection). **b** Quantification of the area of DiI-liposome spread (5 replicate images per 6-12 mice). Data in (**b**) represents mean ± SD, and *p* values were obtained by One-way ANOVA followed by the Tukey's multiple comparison test, $^{**}p<0.01$

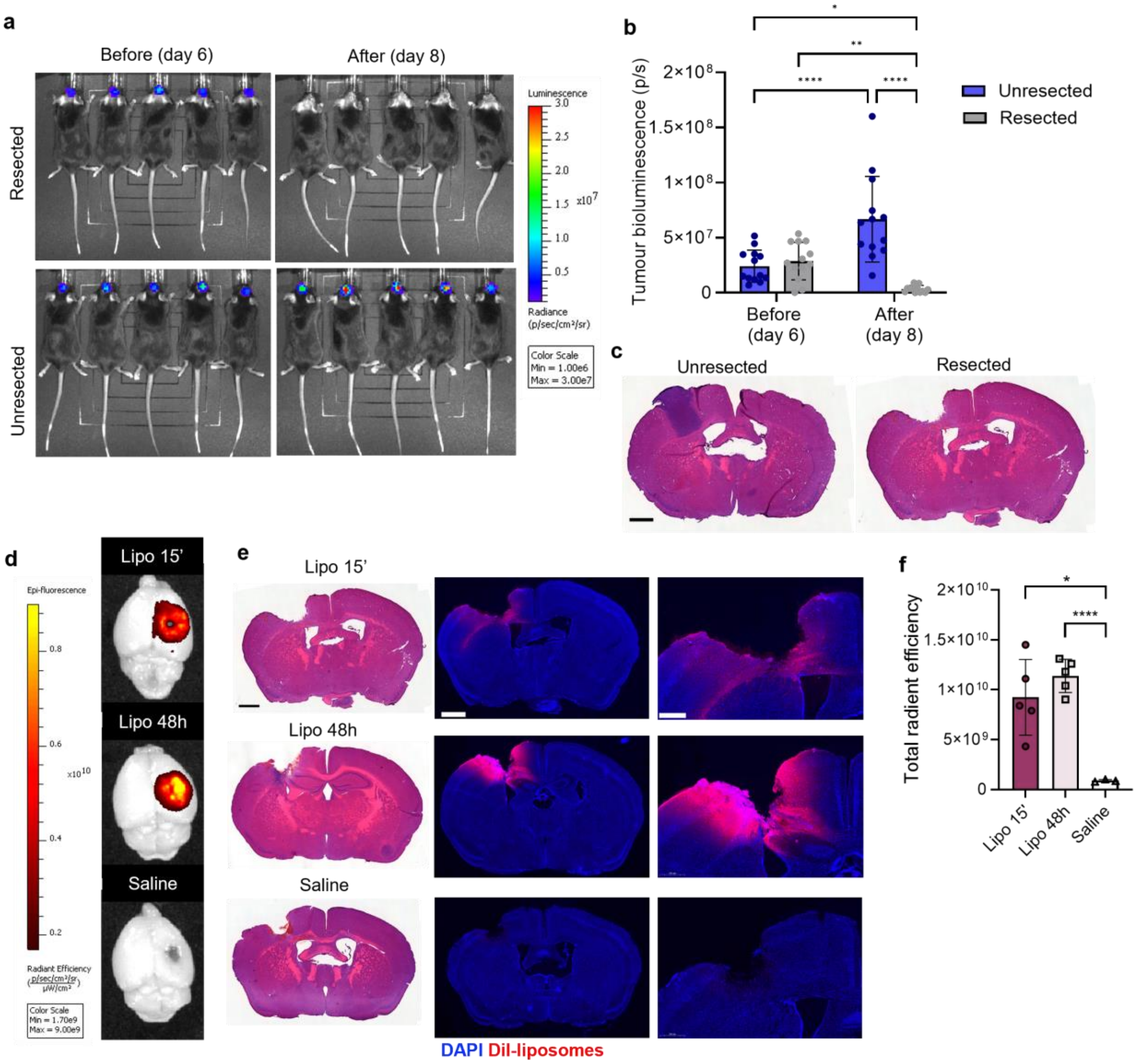


**Supplementary Fig. 6: Validation of DiI-liposome accumulation in a glioma neurosphere (GNS) based glioma resection model. a** Longitudinal *in vivo* imaging of tumor bioluminescence 1 day before (day 6) and 2 days after (day 9) resection of the GNS model. **b** Quantification of total tumor bioluminescence (photons/s) from each animal of the unresected and resected groups overtime ($n$ = 4). **c** H&E histology images of unresected and resected brains on day 2 after resection (days 9 of tumour growth). Representative images from n=4. Scale bar = 1000 µm. **d** Representative *ex vivo* images of perfused brains 24h after liposome injection (at the specified timepoints) showing DiI-liposome distribution in the brain around the surgical site (n=3-5) **e** H&E and associated fluorescence microscopy images of brain sections 24h after i.v. injection of DiI-liposomes or saline and selective accumulation around the resection cavity/margin. Scale bars = 1000 µm. **f** Quantification of liposome fluorescence in the brain expressed as total radiant efficiency ($n$ = 3-5).
Data in (**b,f**) represent mean ± SD, and $p$ values were obtained by **b** two-way ANOVA or **f** one-way ANOVA followed by Tukey's multiple comparison test. *$p$<0.05, **$p$<0.01, ****$p$<0.0001

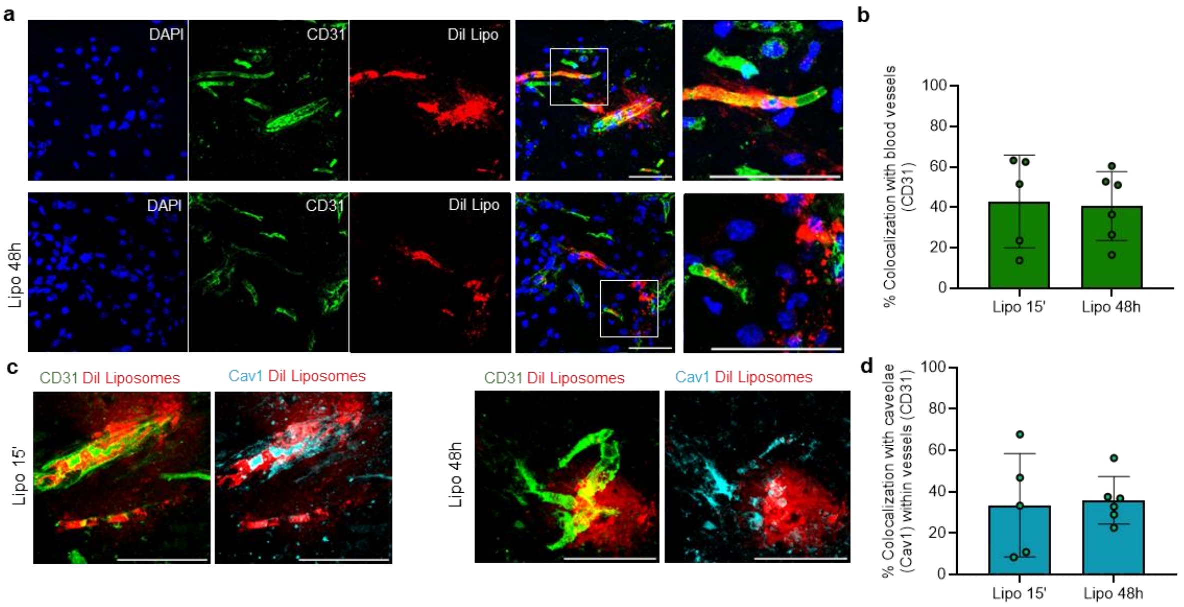


**Supplementary Fig. 7: Liposome interaction with the blood brain barrier. a** Representative confocal micrograph of CD31 (endothelial cells) and DiI-liposomes in mice injected 15' or 48 hours after GBM resection and brains recovered 24 hours after injection with visible liposome extravasation (scale bar = 50 µm). **b** Quantification of percentage signal co-localization of CD31 with DiI-liposomes (4 fields of view per mouse, n= 5-6 mice). **c** Representative confocal micrographs of CD31 (endothelial cells), Cav1 (caveloae) and DiI-liposomes in mice injected 15' or 48 hours after GBM resection and brains recovered 24 hours after injection (scale bar = 50 µm). **d** Quantification of percentage signal co-localization of Cav1 with DiI-liposomes within CD31 vessels (4 fields of view per mouse, n= 5-6 mice).

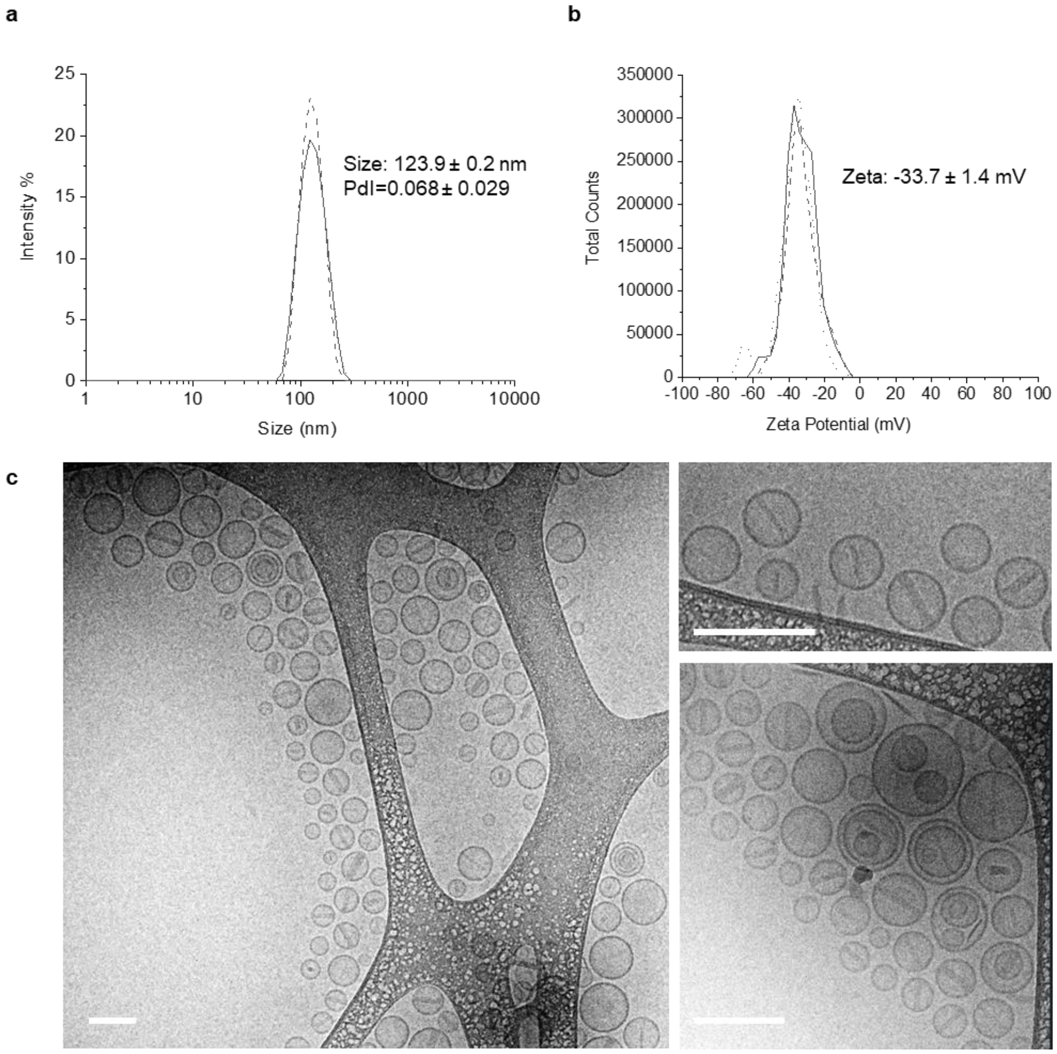


**Supplementary Fig. 8: Doxorubicin-containing liposome (DOX-Lipo) characterization**. **a** Dynamic light scattering (DLS) measurements of size and **b** zeta potential of doxorubicin-containing liposomes. Data is represented as the mean ± standard deviation of 3 measurements. **c** Structural and morphological characterization of doxorubicin-containing liposomes in their native (water) phase by cryoEM. Scale bar = 200 nm.

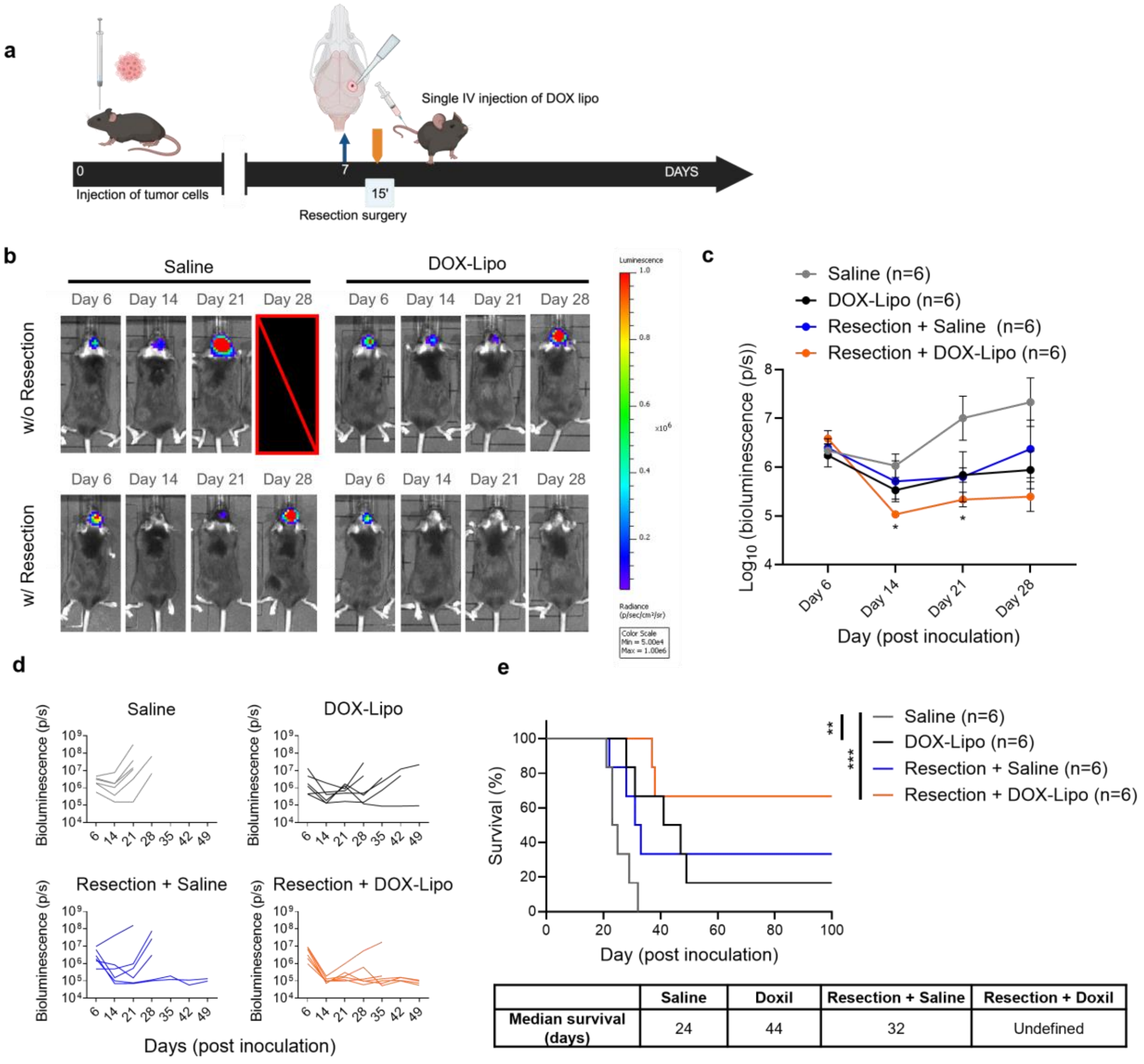


| | Saline | Doxil | Resection + Saline | Resection + Doxil |
|---|---|---|---|---|
| **Median survival (days)** | 24 | 44 | 32 | Undefined |

**Supplementary Fig. 9: Liposomal doxorubicin administration post-resection provides enhanced therapeutic efficacy**. **a** Experimental schematic. Following GBM resection mice received intravenous administration of liposomal doxorubicin (DOX-Lipo) 15’ after surgery. **b** Representative IVIS bioluminescence images on day 6, 14, 21 and 28 post inoculation (days -1, 7, 14 and 21 days post resection) **c** Quantification of tumour bioluminescence (Log10 transformed total flux p/s) within each group (n=6). **d** Quantification of tumour bioluminescence (total flux p/s) of each individual mouse in each group. **e** Survival curves of animals treated with DOX-Lipo or saline vehicle control injected 15’ after surgery with or without tumour resection (n=6). Data in **c** is mean ± SEM. p values were obtained by Two-way ANOVA with Tukey’s multiple comparison test. *p<0.05 relative to saline. *p* values in **d** were obtained with Log-rank (Mantel-Cox) test ***p*<0.01, ***p<0.001.

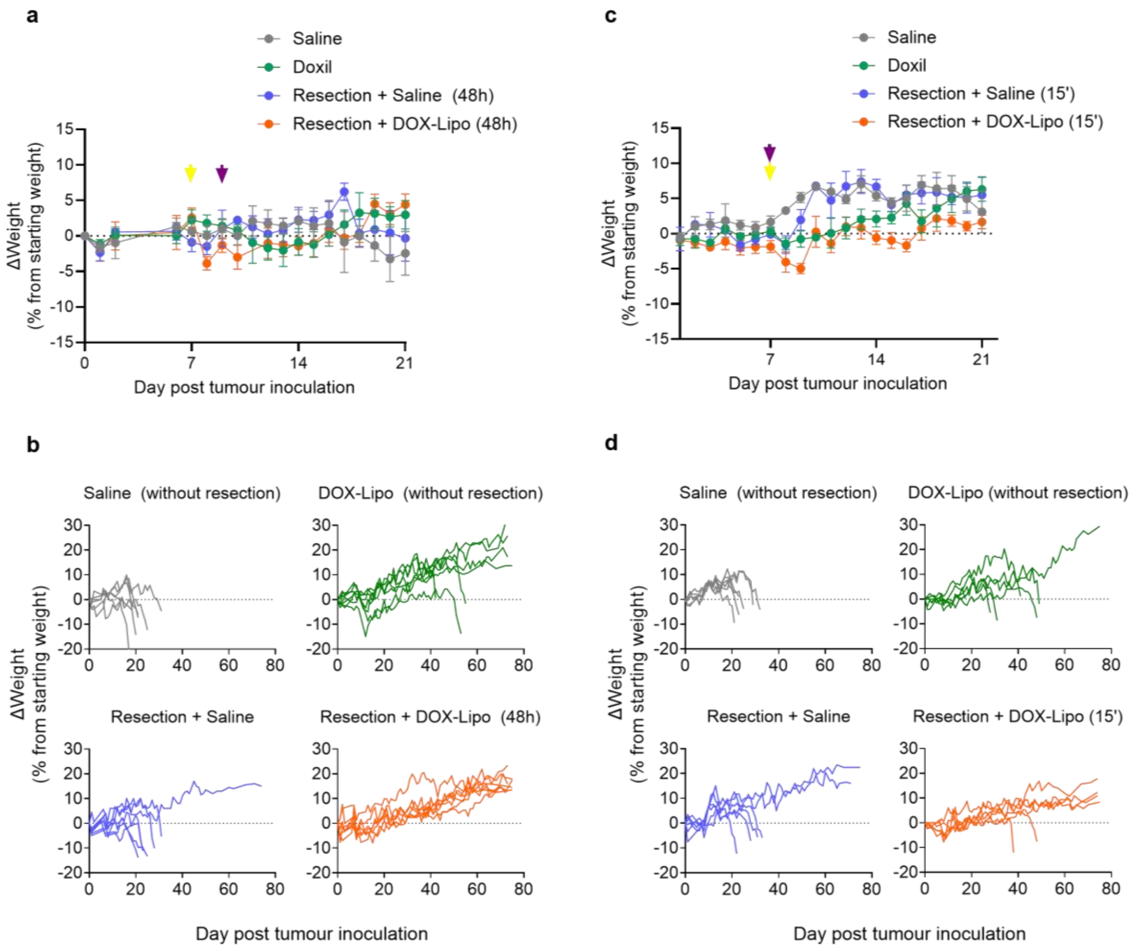


**Supplementary Fig. 10: Animal health under early post-operative Dox-lipo treatment**. Change (**Δ**) in animal weight as a percentage of starting weight present as **a,c** mean ± S.E.M (n=6-7) during acute phase (21 days) and **b,d** individual mice throughout the studies. Yellow/purple arrow in **a**,**c** indicates time of resection and DOX-Lipo injection respectively.
